# Expression noise facilitates the evolution of gene regulation

**DOI:** 10.1101/007237

**Authors:** Luise Wolf, Olin K. Silander, Erik van Nimwegen

**Affiliations:** Biozentrum, University of Basel, and Swiss Institute of Bioinformatics, Basel, Switzerland

## Abstract

In studies of gene regulation, it is often tacitly assumed that the interactions between transcriptional regulators and their target promoters are finely tuned to ensure condition-appropriate gene expression of the targets. However, how natural selection might evolve such precise regulation from an initial state without regulation, is rarely discussed. In addition, the accuracy of gene regulation is affected by noise in gene expression [1]. Expression noise varies greatly across genes [2–5], suggesting that natural selection has affected noise levels, but the role of expression noise in gene regulation is currently poorly understood [6]. Here we present a combination of experimental evidence and theoretical modeling showing that the transmission of expression noise from regulators to their targets can function as a rudimentary form of gene regulation that facilitates the evolution of more finely tuned gene regulation. To assess how natural selection has affected transcriptional noise in *E. coli*, we evolved a large set of synthetic promoters under carefully controlled selective conditions and found, surprisingly, that native *E. coli* promoters show no signs of having been selected for minimizing their noise. Instead, a subset of native promoters, which are characterized by high expression plasticity and high numbers of regulatory inputs, show elevated noise levels. A general theoretical model, which recognizes that target genes are not only affected by the condition-dependent activities of their regulators, but also by the regulators’ noise, explains these observations. Noise transmission from regulators to their targets is favored by selection whenever regulation is imprecise, and may even constitute the main function of coupling a promoter to a regulator. Our theory provides a novel framework for understanding the evolution of gene regulation, demonstrating that in many situations expression noise is not the mere unwanted side-effect of regulatory interactions, but a beneficial function that is key to the evolvability of regulatory interactions.

## Introduction

Studies of gene expression noise in several different model organisms have shown that the promoters of some genes exhibit much more transcriptional noise than others [2–4]. It is unclear, however, how these differences in noise levels have been shaped by natural selection. On the one hand, it can be argued that in each condition there is an optimal expression level for each protein, such that variations away from this optimal level are detrimental to an organism's fitness, implying that selection will act to minimize noise. Indeed, many studies have used circumstantial evidence to suggest that selection generally acts to minimize noise [2, 3, 7–9]. In this interpretation, genes with lowest noise have been most strongly selected against noise, whereas high noise genes have experienced much weaker selection against noise. On the other hand, gene expression noise generates phenotypic diversity between organisms with identical genotypes, and there are well-established theoretical models showing that such phenotypic diversity can be selected for in fluctuating environments [10, 11]. In addition, there is empirical evidence that selection has acted to increase expression noise in some cases [12–15]. It is thus possible that some of the genes with elevated noise may have been selected for phenotypic diversity.

## Results

In order to assess how natural selection has acted on the transcriptional noise of promoters, it is critical to determine what default noise levels would be exhibited by promoters that have not been selected for their noise properties. To address this, we evolved a large set of synthetic *E. coli* promoters *de novo* in the laboratory using an experimental protocol in which promoters were selected on the basis of the mean expression level they conferred, while experiencing virtually no selection on their noise properties (**Fig. 1** and **Supplementary Text**). Starting with an initial library of 100-150 base pair long random sequences, we performed five rounds of mutation and selection, resulting in a genetically diverse collection of functional promoters that conferred expression close to a pre-specified target level (**Fig. 1a-c** and **Suppl. Fig. S1**). We selected a subset of 479 synthetic promoters from the third and fifth rounds, choosing equal numbers of promoters from each of six replicate lineages we evolved (**Fig. 1**; **Methods**). We then used flow cytometry, as described previously [3], to measure the distribution of fluorescence levels per cell for each synthetic promoter, as well as for all native *E. coli* promoters [16].

**Figure 1.**
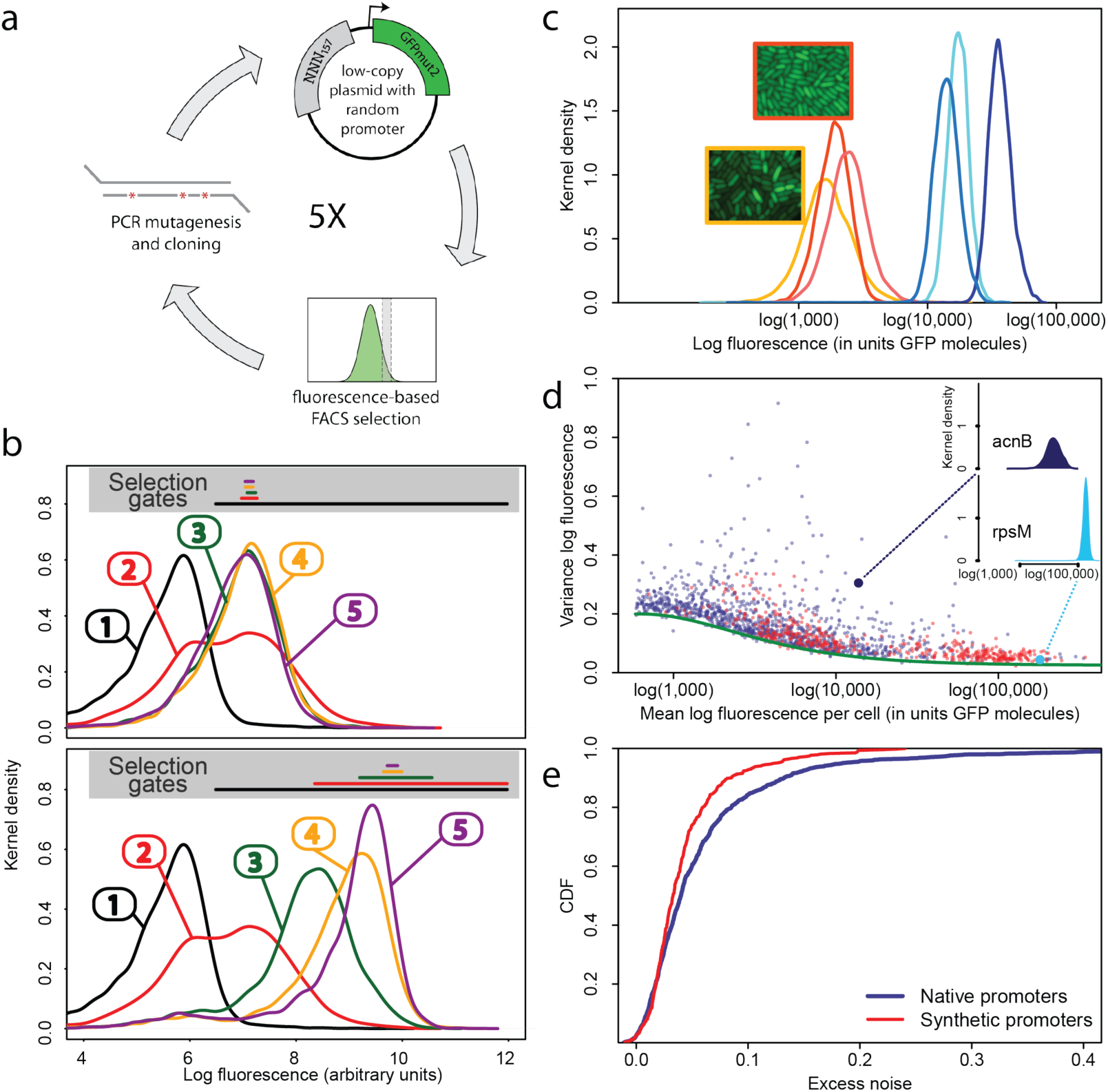
Experimental evolution of functional promoters *de novo*. a: We created an initial library of approximately 10^6^ unique synthetic promoters by cloning random nucleotide sequences, of approximately 100 – 150 base pairs (bp) in length, upstream of a strong ribosomal binding site followed by an open reading frame for GFP as used to quantify the expression of native *E. coli* promoters [16], and transformed this library into a population of cells (**Methods**). We evolved populations of synthetic promoters by performing 5 rounds of selection and mutation on this library. In each round we used fluorescence activated cell sorting (FACS) to select 2 ∗ 10^5^ cells that lie within a gate comprising the 5% of the population closest in fluorescence to a given target level. The plasmids were isolated from the selected cells and PCR mutagenesis was used to introduce new genetic variation into the promoter regions. We then re-cloned the mutated promoters into fresh plasmids, and transformed them into a fresh population of cells. We performed this evolutionary scheme on three replicate populations in which we selected for a target expression level equal to the median expression level (50th percentile) of all native *E. coli* promoters, and three replicate populations in which we selected for a target expression level at the 97.5th percentile of all native promoters (referred to here as medium and high expression levels, respectively). **b:** Changes in the fluorescence distribution for one evolutionary run selecting for medium target expression (top) and one evolutionary run selecting for high target expression (bottom). The curves show the population's expression distributions before selection, with the numbers above each curve indicating the selection round. The colored bars at the top indicate the FACS gates that were used to select cells from the populations at each corresponding round. **c:** Examples of fluorescence distributions for individual clones obtained after five rounds of evolution. Microscopy pictures of two individual clonal promoter populations are shown as insets. **d:** For each native *E. coli* promoter (blue) and synthetic promoter (red) the mean (x-axis) and variance (y-axis) of log-fluorescence intensities across cells were measured using flow cytometry. Fluorescence values are expressed in units of number of GFP molecules. The green curve shows the theoretically predicted minimal variance as a function of mean expression (**Supplementary Text**). The insets show the log-fluorescence distributions for two example promoters (corresponding to the larger dark blue and light blue dots). **e:**, Cumulative distributions of excess noise levels of native (blue) and synthetic (red) promoters.

Observing that the fluorescence distributions across cells were well approximated by log-normal distributions (**Fig. 1c**), we characterized each promoter's distribution by the mean and variance of log-fluorescence, defining the latter as the promoter's noise level (**Fig. 1d**). This definition of noise is equivalent to the square of the coefficient of variation whenever fluctuations are small relative to the mean, which applies to most promoters. Using quantitative Western blotting and qPCR we confirmed that the mean fluorescence levels were directly proportional to GFP molecule numbers and that protein levels were determined primarily by mRNA levels, demonstrating that fluorescence reflected transcriptional activity (**Suppl. Figs S2** and **S3** and **Supplementary Text**). Noise levels were reproducible across biological replicates (**Suppl. Fig. S4**), and noise levels estimated using microscopy were consistent with those measured by flow cytometry (**Suppl. Fig. S5**).

As expected [2,17] we observed a strong relationship between the mean and variance of expression levels of each promoter (**Fig. 1d**). In particular, we observed a strict lower bound on variance as a function of mean expression. This lower bound is well described (**Fig. 1d**, green curve) by a simple model that incorporates background fluorescence, and intrinsic and extrinsic noise components [18] (**Supplementary Text**). We defined the *excess noise* of a promoter as its variance above and beyond this lower bound, allowing us to compare the noise levels of promoters with different means (**Suppl. Fig. S6**). We found, surprisingly, that most of the synthetic promoters exhibited noise levels close to the minimal level exhibited by the native promoters (**Fig. 1d**). Additionally, a substantial fraction of native promoters exhibited excess noise levels significantly greater than the synthetic promoters (**Fig. 1e** and **Suppl. Figs. 6** and **7**). For example, only 26.1% of the synthetic promoters exhibited excess noise above 0.05, compared to 41.6% of the native *E. coli* promoters (*p* < 7.7 10*^−^*^10^, hypergeometric test). Given that the synthetic promoters were evolved from random sequence fragments, and had not been selected on their noise properties (**Supplementary Text**), we concluded that constitutively expressed *E. coli* promoters exhibit low excess noise levels by default. Importantly, this implies that the native promoters with elevated excess noise must have experienced selective pressures that caused them to increase their noise.

To understand how selection might have acted to increase noise, we first investigated whether excess noise was associated with other characteristics of the promoters. Previous studies in *S. cerevisiae* have shown that promoters with high noise tend to also show high expression plasticity, i.e. large changes in mean expression level across environments [2]. Although we did not clearly observe this association in data from our previous study [3], a recent re-analysis of this data did uncover a significant association between expression plasticity and noise [19], which we confirmed using our present data (**Fig. 2a**). In addition, we found that there is an equally strong relationship between excess noise and the number of regulators known to target the promoter [20] (**Fig. 2b**). In particular, whereas the excess noise levels of promoters without known regulatory inputs are very similar to those of our synthetic promoters, promoters with one or more regulatory inputs have clearly elevated noise levels (**Fig. 2c**).

**Figure 2.**
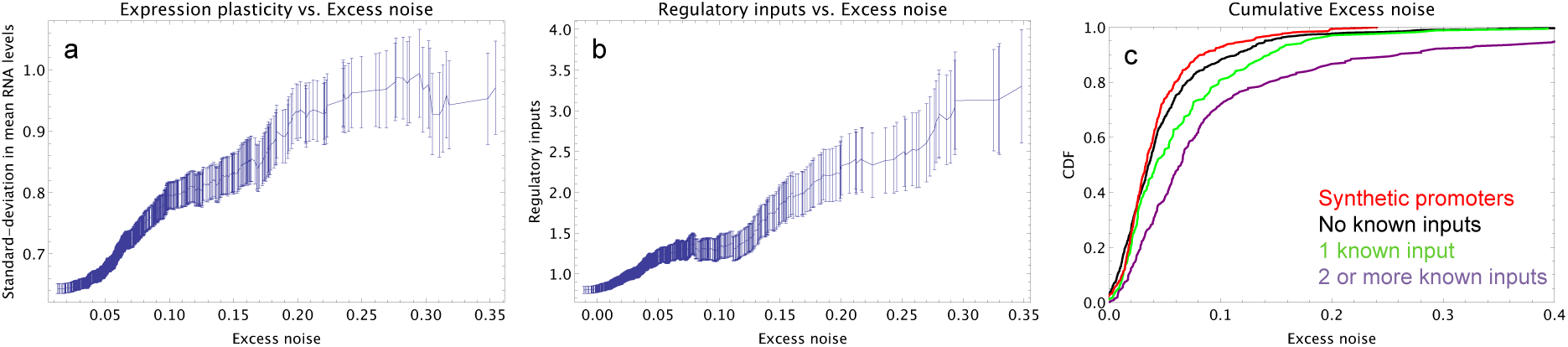
Promoters with elevated noise exhibit high expression plasticity and large numbers of regulatory inputs. a: Native promoters were sorted by their excess noise *x* and, as a function of a cut-off on *x* (horizontal axis), we calculated the mean and standard-error (vertical axis) of the variation in mRNA levels across different experimental conditions (data from http://genexpdb.ou.edu/) of all promoters with excess noise larger than *x*. **b:** Promoters were sorted by excess noise *x* as in panel a, and mean and standard-error of the number of known regulatory inputs (vertical axis, data from RegulonDB [20]) for promoters with excess noise larger than *x* is shown. **c:** Cumulative distributions of excess noise levels of synthetic promoters (red) and native promoters without known regulatory inputs (black), with one known regulatory input (green), and with two or more known regulatory inputs (purple).

We next considered what the origin of this general association between noise and regulation could be. It is important to recognize that, when a promoter couples to a transcription regulator by evolving cognate binding sites, the expression of the associated gene will be affected in two separate ways. First, the gene's mean expression will become correlated with the activity of the regulator in a condition-specific manner. Second, in addition to this ‘condition-response’ effect, the noise in the expression or activity of the regulator will be transmitted to the target gene. This ‘noise-transmission’ effect will cause an increase in expression noise of the target [21]. Based on this noise-transmission effect, and in analogy with fluctuation-dissipation theorems from physics, it has been proposed that elevated expression noise is simply an unwanted but unavoidable side-effect of regulation [22].

However, there is no reason to assume that the condition-response and noise-transmission effects must always be in selective conflict with each other. Several theoretical treatments have shown that phenotypic variability may be selectively beneficial when environments change in ways that cannot be accurately sensed or are too rapid for organisms to respond [10, 11, 23]. Although such theoretical studies are typically less concerned with the mechanisms by which such increased noise could be genetically encoded, the noise-transmission effect is one obvious candidate mechanism. It would thus seem that, at least in some situations, the condition-response and noise-transmission effects could act in concert. To quantify how selection might act on the combination of these two effects, we developed a general model that considers a gene whose optimal expression levels vary across conditions, and calculated how the condition-response and noise-transmission effects of coupling to a given regulator conspire to affect fitness (**Supplementary Text**). Although our model applies very generally (**Supplementary Text**), we illustrate it here using a simple scenario (**Fig. 3**).

**Figure 3.**
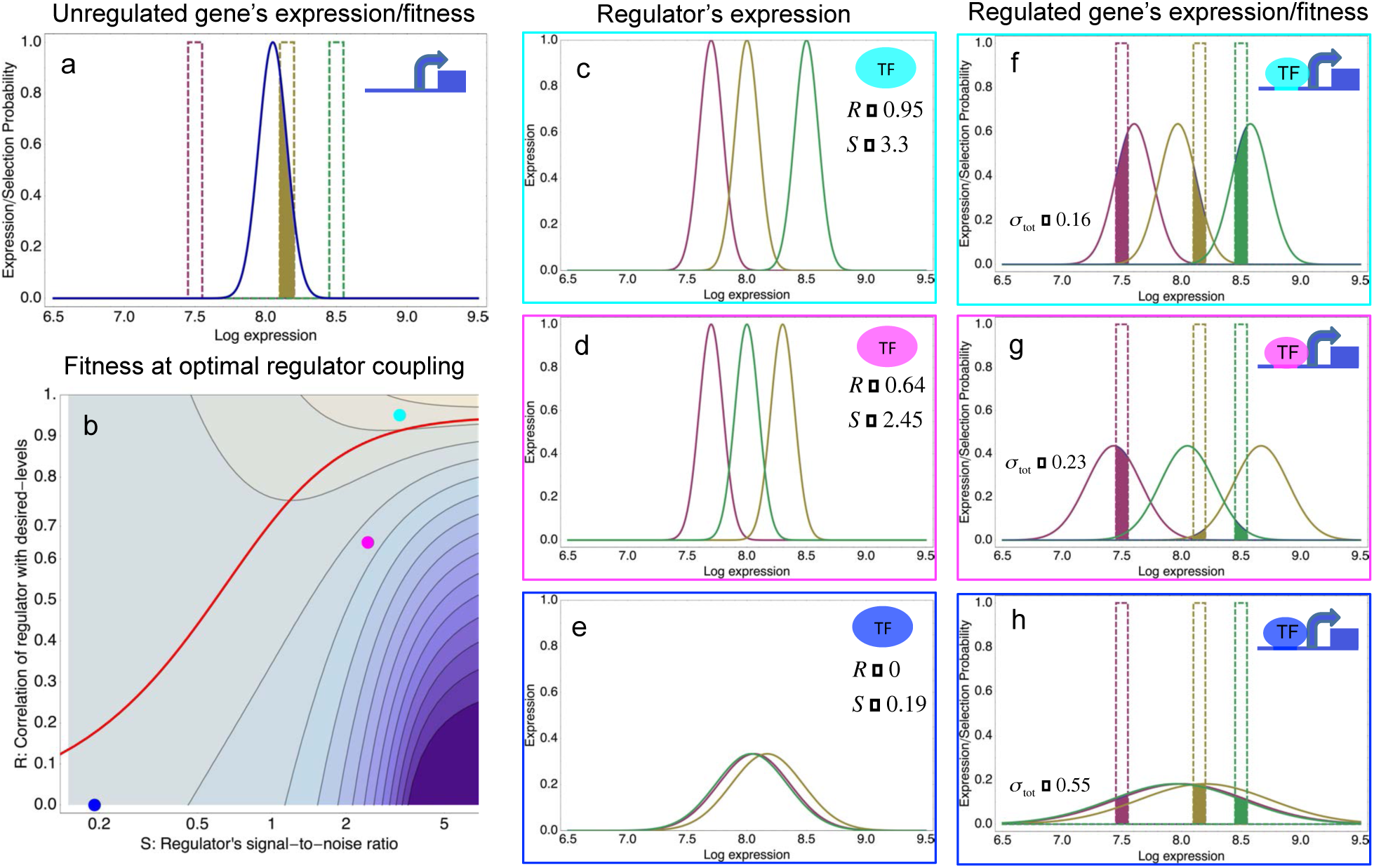
A model of the evolution of gene expression regulation in a variable environment. a: Expression distribution of an unregulated promoter (blue curve) and selected expression ranges in 3 different environments, i.e. the red, gold, and green dashed curves show fitness as a function of expression level in these environments. Although our model applies more generally, for simplicity we here visualize selection as truncation selection (i.e. a rectangular fitness function). The fitness of the promoter in the gold environment is proportional to the shaded area. **b:** Contour plot of the log-fitness of a promoter that is optimally coupled to a transcription factor (TF) with signal-to-noise ratio *S* and correlation *R*. Contours run from 0.5 at the top right to −7.5 at the bottom right. The three colored dots correspond to the TFs illustrated in panels **c-h**. The red curve shows optimal S as a function of *R*. **c-e:** Each panel shows the expression distributions of an example TF across the 3 environments (red, gold, and green curves). The corresponding values of correlation *R* and signal-to-noise *S* are indicated in each panel. **f-h:** Each panel shows the expression distributions across the 3 environments for a promoter that is optimally coupled to the TF indicated in the inset. The shaded areas correspond to the fitness in each environment. The total noise levels of the regulated promoters are also indicated in each panel. The unregulated promoter has total noise *σ*_tot_= 0.1.

The expression of an unregulated promoter is characterized by a distribution with a given mean and variance (**Fig. 3a**, blue curve). We assume that the organism experiences different environments and that, in each environment, cells with expression levels within a certain range are selected. In the simple scenario of **Fig. 3** there are 3 environments (red, gold, and green), with the green environment requiring up-regulation of the expression and the red environment requiring down-regulation of the expression (**Fig. 3a**). The fitness in each environment corresponds to the fraction of cells with expression levels within the selected range, i.e. the unregulated promoter has reasonably high fitness in the gold environment but very low fitness in the green and red environments. Since the organisms experience all 3 environments, a poor overlap between the expression distribution and the selected range in any one environment leads to low overall fitness.

To improve fitness, a promoter may evolve binding sites for an existing regulator, such that its expression becomes dependent on the activity of this regulator, which will generally vary across environments. Our modeling shows that the resulting fitness depends only on 2 effective parameters of the regulator: The correlation *R* between the condition-dependent expression levels (or, more generally, activities) of the regulator and the desired levels of the promoter, and the signal-to-noise parameter *S* that characterizes the accuracy of the regulator's expression.

As intuitively expected, the highest fitness is obtained when coupling to an accurate regulator with high signal-to-noise *S*, i.e. whose activities correlate precisely with the desired expression levels (cyan dot in **Fig. 3b** and **Fig. 3c,f**). The resulting expression distributions of the promoter accurately track the desired levels, with only moderately increased noise in the promoter's expression (**Fig. 3f**). However, regulators that track the desired expression levels of the promoter with such high accuracy may often not be available. Interestingly, coupling to a noisy regulator whose activity is entirely uncorrelated with the desired expression levels (blue dot in **Fig. 3b** and **Fig. 3e,h**) also substantially increases fitness. In this regime, the increased fitness results exclusively from the noise-transmission mechanism. Surprisingly, coupling to the uncorrelated noisy regulator (blue dot in **Fig. 3b** and **Fig. 3e,h**) outperforms coupling to a moderately correlated regulator (magenta dot in **Fig. 3b** and **Fig. 3d,g**). This is due to the fact that the magenta regulator is not noisy enough given its correlation *R*, i.e. lowering *S* for this TF would result in an increase of the promoter's noise, and this would lead to an increase in fitness in the green and gold conditions (see **Fig. 3g**). This illustrates that regulators may be under selection to become noisy themselves and the red curve in **Fig. 3b** shows the optimal signal-to-noise *S* of a regulator as a function of its correlation *R*. Whereas to the right of this curve the noise-transmission is too large, and too small to the left of it, along the curve the condition-response and noise-transmission effects are optimally acting in concert. This clarifies how accurate gene regulation can evolve smoothly, starting from a noisy regulator with low *R* and *S* whose benefits come entirely from the noise-transmission, by increasing both *R* and *S* in small steps, until reaching highly accurate regulation with high *R* and *S* for which the condition-response effect dominates.

Our model also predicts how the final noise of a promoter depends on the variance in its desired expression levels (**Supplementary Text**). In particular, assuming the best available regulator in the genome has a given correlation *R* with the desired levels, there will be a critical variance such that below this variance the final noise will be equal to the noise of the unregulated promoter, and above this critical variance the final noise of the promoter will be proportional to (**Suppl. Fig. S8**). That is, our model explains the observation that expression noise increases with expression plasticity. Similarly, in our simple model the increase in expression noise is directly due to coupling to regulators, such that our model also explains the observed general association between expression noise and regulatory inputs.

## Discussion

Because genotype-phenotype relationships for complex phenotypic traits are poorly understood, it is often difficult to assess how observable variation in a particular trait has been affected by natural selection. Here we have shown that by comparing naturally observed variation in a particular trait with variation observed in synthetic systems that were evolved under well-controlled selective conditions, definite inferences can be made about the selection pressures that have acted on the natural systems. In particular, by evolving synthetic *E. coli de novo* using a procedure in which promoters are strongly selected on their mean expression and not on their expression noise, we have shown that native promoters must have experienced selective pressures that increased their noise levels. To account for this, we have proposed a theoretical model that provides a simple mechanistic framework for understanding how selection can act to couple transcriptional regulators and target genes, and which quantifies the parameter regimes in which we expect promoters to exhibit high levels of noise. This framework vastly expands the evolutionary conditions under which novel regulatory interactions will be selected for; instead of assuming that the regulators and their targets must evolve in a tightly coordinated fashion, the model shows that genes may often benefit from coupling to regulators whose activities do not correlate with the gene's expression requirements at all. In particular, the condition-response and noise-transmission effects of coupling to a regulator, rather than being in conflict with each other, may often act in concert. Finally, our model shows quite generally that unless regulation is very precise, regulatory interactions that act to increase noise are beneficial. Thus, high levels of expression noise can be expected whenever the accuracy of regulation is limited.

## Methods

### Ab initio promoter library construction from random sequences

We obtained chemically synthesized nucleotide sequences of random nucleotides 200 bp in length (Purimex, Germany). Each sequence had defined 5’ and 3’ ends to allow PCR amplification. Within these constant regions, restriction sites for BamHI and XhoI were present. The intervening sequence was made up of 157 bp of random nucleotides (5’-CCTTTCGTCTTCACCTCGAG-(N157)-GGGATCCTCTGGATGTAAGAAGG-3’). However, as coupling of base pairs during oligonucleotide synthesis is not always successful and strand breaks can frequently occur in long oligonucleotides, many oligonucleotides were shorter than 200 bp in length. We used PCR to generate double stranded DNA from the single stranded oligonucleotides using forward and reverse primers matching the defined 5’ and 3’ ends. We gel-purified the double-stranded PCR product and double-digested it using BamHI and XhoI. After column-purification, sequences were ligated into a version of the low-copy plasmid pUA66, which contains a gfpmut2 open reading frame downstream of a strong ribosomal binding site [16]. The vector was modified to remove a weak *σ*70 binding site present 24 bp upstream of the GFP open reading frame (two point mutations, A→G and T→G, were introduced, changing the putative *σ*70 binding site from TAGATT to TGGATG, with the consensus *σ*70 binding site being TATAAT). The ligation was performed using T4 DNA ligase (NEB) at 16°C for 24 hours. The ligation product was then column purified and electroporated into *E. coli* DH10B cells. This protocol resulted in extremely high transformation yields (approximately 10^6^ individual clones per transformation).

### Selection on expression level using flow cytometry

Cultures of transformed cells were regenerated for one hour in 1 mL SOC medium (Super Optimal Broth supplemented with 20 *mM* glucose) and afterwards 1 *mL* SOC containing 50 µ*g/ml* kanamycin was added for overnight growth, ensuring that only cells containing the plasmid could grow. These cultures were then diluted 500-fold (approximately 5 * 10^6^ cells in total) into M9 minimal media supplemented with 0.2% glucose and grown for 2.5 hours with shaking at 200 rpm. The distribution of GFP fluorescence levels was measured for each culture using fluorescence activated cell sorting (FACS) in a FACSAria IIIu (BD Biosciences), with excitation at 488 nm and a 513/17 nm bandpass filter used for emission.

We used this distribution of fluorescence values to designate a selection gate. The position of the gate was determined by measuring the mean fluorescence of two reference promoters [16]: *gyrB* which exhibits a mean expression level that is at the 50th percentile all *E. coli* promoters; and *rpmB*, which exhibits a mean expression level that is at the 97.5th percentile of all *E. coli* promoters [3]. For each of these reference genes, the mean fluorescence level was measured, and a selection gate was constructed, centered on this mean expression level, such that 5% of all clones in the population fell within the gate. For each round of selection, we sorted 200′000 cells contained within this gate. Sorted cells were then transferred to 4 *mL* Luria Broth (LB) media (containing 50 µ*g/ml* Kanamycin) and grown overnight. These cultures were stored supplemented with 7.5% glycerol at −80°C for subsequent analysis.

For each expression level (i.e. reference gene), we evolved three replicate populations. We refer to these as the medium expressers (those promoters selected based on the *gyrB* reference gate) and high expressers (those promoters selected based on the *rpmB* reference gate).

### PCR mutagenesis

Following FACS-based selection on fluorescence, we introduced novel genetic variation into the populations using PCR mutagenesis. We first re-grew the cells overnight and used this culture to prepare plasmid DNA. We amplified the promoter sequences from these plasmids using the GeneMorph II Random Mutagenesis Kit (Stratagene) with the primers referred to previously that matched the defined regions of the promoters. We used 0.01 ng of DNA as starting material and 35 cycles for amplification. This resulted in a mutation rate of around 0.01 per bp (such that we expect that in 200 bp, 95% of the promoters will contain between zero and four mutations). These PCR products were then digested with XhoI and BamHI, ligated back into the vector, and again transformed into DH10B cells. We repeated this entire process (selection, PCR mutagenesis, and transformation) five times in total. At this point, the plasmid libraries of synthetic promoters were isolated and transformed into *E. coli* K12 MG1655 for comparison to a library of native *E. coli* promoters (see below).

### Quantification of fluorescence

To quantify fluorescence on a single-cell level, we used flow cytometry with a FACSCanto II (BD Biosciences), with excitation at 488nm and a 513/17 nm band-pass filter used for emission. We collected data for at least 50′000 events. We then gated this data as outlined in [3], identifying approximately 5′000 cells most similar in FSC and SSC. We then calculated the mean and variance in log-fluorescence using these cells, using a Bayesian procedure that accounts for outliers (**Supplementary Text**). We randomly selected 479 promoters from the evolved set (72 medium expressers and 72 high expressers after 3 rounds of selection; 168 medium expressers and 167 high expressers after 5 rounds of selection) and quantified mean and variance in fluorescence. We used the same measurement procedures to calculate mean and variance for all promoters contained in a library of *E. coli* promoters also placed upstream of the gfpmut2 open reading frame on the pUA66 plasmid [16]. We refer to the promoters from this library as native *E. coli* promoters. For 288 promoters, we quantified fluorescence in three independent cultures and found that both mean and variance in expression were reproducible across replicate biological experiments (**Suppl. Fig. S4**). Additionally, we sequenced 378 sequences from our set of 479 promoter sequences, which showed that even after five rounds of selection, the promoters were quite diverse (**Suppl. Fig. S1**). To confirm the sensitivity and accuracy of the FACS measurements, we selected ten promoters and used fluorescence microscopy to measure their mean and variance in fluorescence. The cells were grown in the same conditions described above, placed on 1% agarose pad, and images were obtained using a CoolSNAP HQ CCD camera (Photometrics) connected to a DeltaVision Core microscope (Applied Precision) with a UPlanSApo 100X/1.40 oil objective (Olympus). Image-processing was done in soft-WoRx v3.3.6 (Applied Precision) and fluorescence values were extracted based on DIC-image mediated cell detection in MicrobeTracker Suite [24]. For each cell, we calculated fluorescence per cell volume by summing all pixel values and dividing by the volume of the cell as estimated by MicrobeTracker. Cells undergo substantial phenotypic changes when they are put on agar, including changes in the distribution of cell sizes. Consequently, it is problematic to compare absolute variance measurements directly between FACS and microscope. We therefore compared the relative noise levels of different promoters. The 10 selected native promoters consist of 5 pairs with almost identical mean expression values (as measured by the FACS) but with noise levels that vary by different amounts. For each of the 5 pairs, we calculated the ratio of the noise levels of the higher and the lower noise promoter as measured by both the FACS and the microscope. As shown in **Suppl. Fig. S5**, with the exception of one pair of promoters that showed almost equal noise levels in the FACS but a 50% difference in noise in the microscope, all other pairs showed good correlation of the relative noise levels in the FACS and in microscope, confirming that relative noise levels are similar in FACS and microscope measurements.

### Quantitative Western analysis

To determine the correspondence between fluorescence intensities and absolute GFP numbers per cell, eight individual promoter clones were grown in three biological replicates using the same media conditions as in the experimental evolution. The cells were then re-suspended in SDS sample buffer, heated for 5 minutes at 95°C, and proteins were resolved by 12% SDS-PAGE. Quantification was done by loading a standard curve consisting of 10, 25, 50, 75, and 100 nanograms of GFP (Clonetech, #632373). Proteins were transferred to a Hybond ECL membrane (GE Healthcare, Life Sciences), which was then blocked in TNT (20 mM Tris pH 7.5, 150 mM NaCl, 0.05% Tween 20) with 1% BSA and 1% milk powder. Detection was performed with the ECL system after incubation with rabbit anti-GFP and polyclonal pig anti-rabbit. Western intensities for each sample were extracted using ImageJ (**Suppl. Fig. S2**). The number of cells loaded was estimated by calculating the relationship between OD600 and CFU counts. Details of the data analysis procedures are in the **Supplementary Text**.

### Correlating protein and RNA levels per cell by quantitative PCR

Native and evolved single-promoter populations were grown in three biological replicates by diluting overnight LB cultures 500-fold into M9 media supplemented with glucose. These cultures were grown for 2.5 hours, stabilized with an equal volume of RNA Later (Sigma-Aldrich) and RNA was extracted using the Total RNA Purification 96-Well Kit (Norgen Biotek Corp.) with on-column DNAse I digestion. Reverse transcription was done using random hexamers and qPCR with TaqMan probes and performed by Eurofins Medigenomix GmbH (Germany). Three technical replicates were performed. The efficiency of the primers and probes used were validated in a dilution series. Relative RNA levels per cell were obtained by normalizing to the reference gene *ihfB* using a Bayesian procedure for integrating data from the replicates and accounting for failed measurements (**Supplementary Text**). The primers and probes used were: GFP forward primer: 5′-CCTGTCCTTTTACCAGACAA-3′; GFP reverse primer: 5′-GTGGTCTCTCTTTT-CGTTGGGAT-3′; GFP probe: 5′-TACCTGTCCACACAATCTGCCCTTTCG-3′, ihfB forward primer: 5′-GTTTCGGCAGTTTCTCTTTG-3′, ihfB reverse primer: 5′-ATCGCCAGTCTTCGGATTA-3′, ihfB probe: 5′-ACTACCGCGCACCACGTACCGGA-3′).

### Minimal variance as a function of mean expression and excess noise

In a simple model of gene expression in which there are constant rates of transcription, translation, mRNA decay, and protein decay, the probability distribution for the number of proteins per cell is a negative binomial with variance proportional to the mean 〈*n*〉: var(*n*) = (*b* + 1)〈*n*〉, where the constant *b* is the ratio between the mRNA translation rate and the mRNA decay rate, which is often referred to as ‘burst size’ [25]. However, in general there are also cell-to-cell fluctuations in the transcription, translation, and decay rates, which are proportional to these rates themselves. These fluctuations lead to an additional term in the variance var(*n*) which is proportional to the square of the mean: 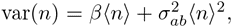 where *β* is a renormalized burst size and 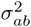 is the relative variance of the product of transcription, translation, and decay rates across cells (**Supplementary Text**).

The total fluorescence in a cell (measured in units equivalent to number of GFP proteins) *n*_meas_ can then generally be written as: 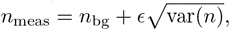 where *n*_bg_ is background fluorescence and *∈* is a fluctuating quantity with mean zero and variance one. Assuming that the fluctuations are small relative to the mean, we then find for the variance of the logarithm of *n*_meas_:

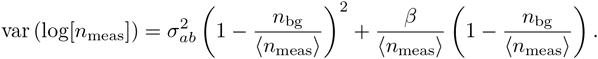

We fit this functional form to the minimum variance var (log[*n*_meas_]) as a function of the mean, with 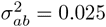 and *β* = 450. We defined the excess variance as the difference between the measured variance and this fitted minimal variance. A more detailed derivation is given in the **Supplementary Text**.

### The FACS selection function

By comparing the distributions of the population's expression levels before and after rounds of selection (without intervening mutation of the promoters), we found that the probability that a cell with expression level *x* is selected by the FACS is well-approximated as 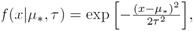 with *µ* the desired expression level and *τ* the width of the selection window. For the last 3 rounds of selection for medium expression, we estimated *τ* ≈ 0.03 and *µ_∗_* fluctuated slightly around an average value of *µ_∗_* ≈ 8.1.

With this selection function, a promoter genotype that exhibits a distribution of expression values with mean *µ* and standard-deviation *σ* has a fitness (fraction of cells selected in the FACS) of

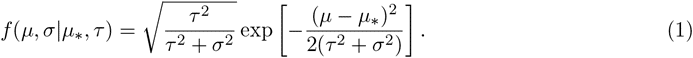

This estimated fitness function indicated that the fitness of promoter genotypes strongly depends on their mean *µ* and is almost independent of their excess noise. In addition, applying additional rounds of selection of varying strengths to the population of evolved promoters did not systematically alter their distribution of excess noise levels. Details of the analysis of the FACS selection are in the **Supplementary Text**.

### Model for the evolution of gene regulation in a fluctuating environment

Although the model we present can be extended to include the evolution of gene regulation for multiple genes, for simplicity we focused on the evolution of a single gene and its promoter. We assumed that the population experienced a sequence of different environments and that, in each environment, the fitness of each organism is a function of its gene expression level. We characterized the fitness function in each environment by two parameters: the desired level *µ_e_*that maximizes the fitness and a parameter *τ* that quantifies how quickly fitness falls away from this optimum and, for simplicity and analytical tractability, we assumed a Gaussian form: 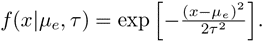 Note that this is the same form as the FACS selection function. Consequently, the fitness *f*(*µ*, *σ|µ_e_*, *τ*) of a promoter with mean *µ* and variance *σ*^2^ is given by equation (1) as well, with *µ_e_* replacing *µ_∗_*.

The total number of offspring that a promoter will leave behind after experiencing all environments is given by the product of its fitness in each of the environments. Equivalently, the log-fitness of a promoter is proportional to its average log-fitness across all environments. We then find for the log-fitness:

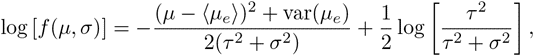

where 〈*µ_e_*〉 is the average of the desired expression levels across environments, and var(*µ_e_*) is the variance in the desired expression levels across conditions. If we do not consider gene regulation, but simply optimize the promoter's mean expression and noise level, then we find optimal log-fitness occurs when *µ* = 〈*µ_e_*〉 and *σ*^2^ = 0 (when var(*µ_e_*) *< τ* ^2^) or *σ*^2^ = var(*µ_e_*) − *τ*^2^ otherwise. That is, when the desired expression level varies more than the width of the selection window, noise is increased so as to ensure the distribution overlaps the desired levels across all conditions. This result is equivalent to previous results on the evolution of phenotypic diversity in fluctuating environments [10].

To increase fitness, a promoter can evolve to become regulated by one of the regulators existing in the genome. Instead of having a constant mean expression *µ*, the promoter's mean expression will then become a function of the environment *e*: *µ*(*e*) = *µ* + *cr_e_*, where *r_e_*is the mean expression (or more generally regulatory activity) of the regulator in environment *e*, and *c* is the coupling strength. Since any gene will have some variability in its expression, we assumed that the actual expression/activity of the regulator in environment *e* is Gaussian distributed with a variance 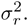 Consequently, when coupled to the regulator, the promoter's total expression variance will become 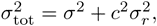 and the log-fitness of the promoter becomes:

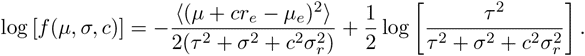

Assuming that the basal expression *µ* is optimized to maximize log-fitness, i.e. *µ* = 〈*µ_e_*〉 − *c〈r_e_*〉, this log-fitness can be rewritten as:

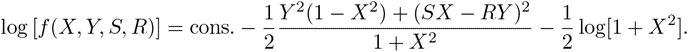

where *X* measures the coupling strength 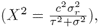 *Y* is the expression mismatch that measures how much the desired expression level varies across environments 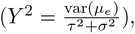 *S* is the signal-to-noise of the regulator 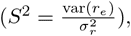 and *R* is the Pearson correlation between the desired expression levels *µ_e_* and the activity levels *r_e_* of the regulator. Additional details on this derivation and analysis of the behavior of the fitness function as a function of its parameters are given in the **Supplementary Text**.

### Analysis of excess noise against gene expression variation and regulatory inputs

We re-annotated the promoter fragments of [16] by mapping the published primer pairs to the *E. coli* K12 MG1655 genome. Of the 1816 promoter fragments, 1718 could be unambiguously associated with a gene that was immediately downstream, and the 1718 promoter fragments were associated with 1137 different downstream genes (for some genes, there were multiple or repeated upstream promoter fragments). We used the operon annotations of RegulonDB [20] to extract, for each promoter, the set of additional downstream genes that are part of the same operon as the first downstream gene. We obtained known regulatory interactions between transcription factors and genes from RegulonDB and counted, for each *E. coli* gene, the number of transcription factors known to regulate the gene. We defined the number of regulatory inputs of a promoter to equal the average of the number of inputs for all genes in the operon downstream of the promoter. We sorted promoters by their excess noise and, as a function of a cut-off on excess noise level, calculated the mean and standard-error of the number of regulatory inputs for all promoters with excess noise level above the cut-off. We obtained genome-wide gene expression measurements from the Gene Expression Database (http://genexpdb.ou.edu/). For each *E. coli* gene, we obtained 240 log fold-change values *x* corresponding to the logarithm of the expression ratio of the gene in a perturbed and a reference condition. We defined the variance in expression of a gene as the average of *x*^2^ across the 240 experiments. We again sorted promoters by their excess noise and, as a function of a cut-off on excess noise level, calculated the mean and standard error of gene expression variances for all promoters with excess noise above the cut-off.

## Acknowledgments

We thank D. Blank for performing the Western Blots, D. Bumann for discussions, B. Claudi and J. Zankl for assistance with flow cytometry, and S. Abel and I. de Jong for assistance with microscopy.

## Contributions

OS and EvN designed the study. Experiments were designed by OS and LW and performed by LW. LW, OS, and EvN analyzed the data, and EvN developed the theoretical model. LW, OS, and EvN wrote the paper.

## Supplementary Figures

**Figure S1.**
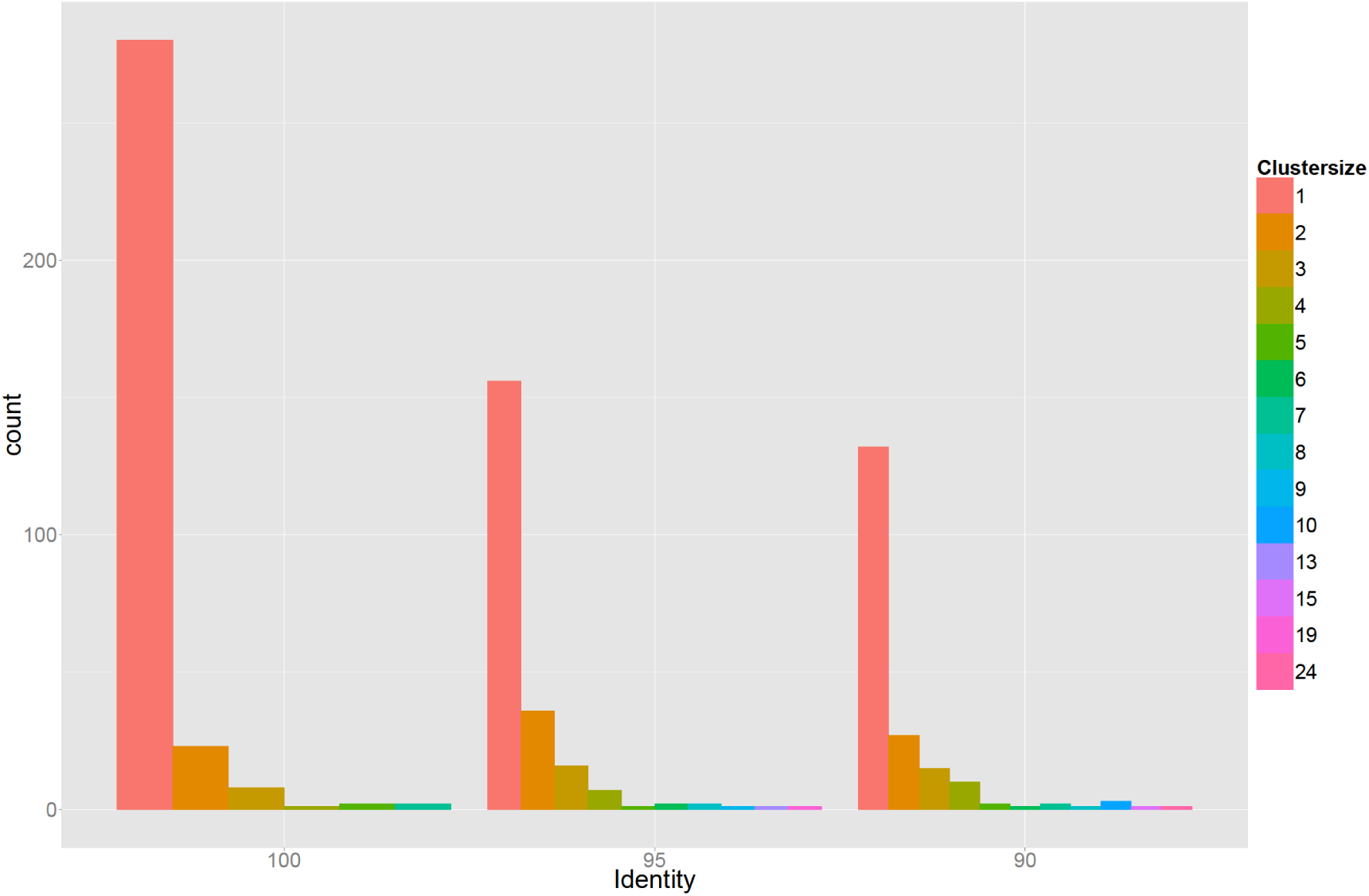
Genetic diversity of 378 sequenced promoters, which were extracted from randomly selected clones from the populations that were obtained after 3 and 5 rounds of selection. Sequences were clustered using single-linkage based on 100, 95, or 90 percent sequence identity (left, middle, and right panel) and the bar plots show the corresponding histograms of cluster sizes. The results indicate that the promoters in the populations at the third and fifth rounds are highly diverse, deriving from many different initial random sequences in the initial library.

**Figure S2.**
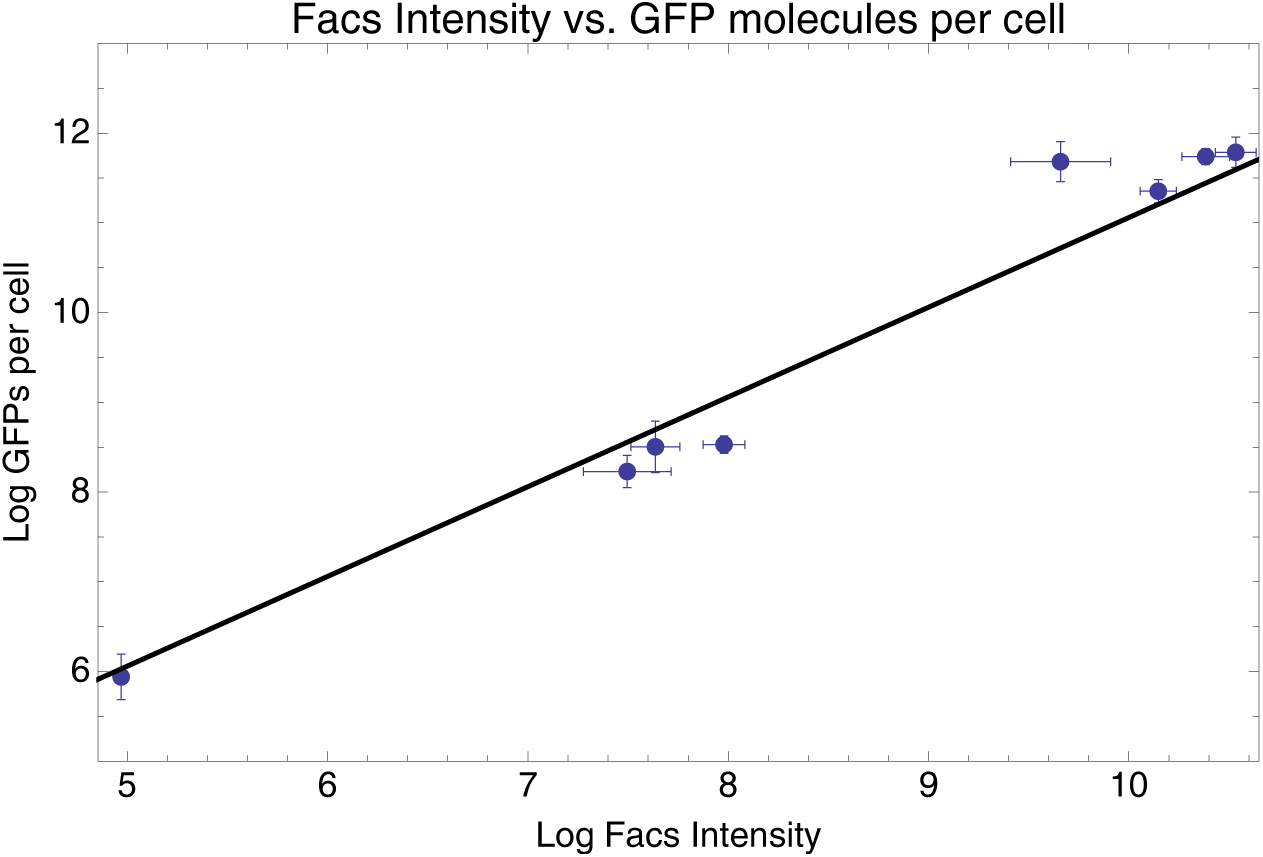
Mean log-fluoresence intensities as measured by FACS (horizontal axis) against estimated log GFP molecules per cell (vertical axis) as estimated from quantitative Westerns (see **Supplementary Text**) for 8 selected promoters. Error-bars were estimated from 3 replicates for the FACS measurements and 6 replicates for the GFP levels. The straight line shows the fit *y* = *x* + 1.06, which is equivalent to: GFP molecules per cell = 2.88 * mean FACS intensity.

**Figure S3.**
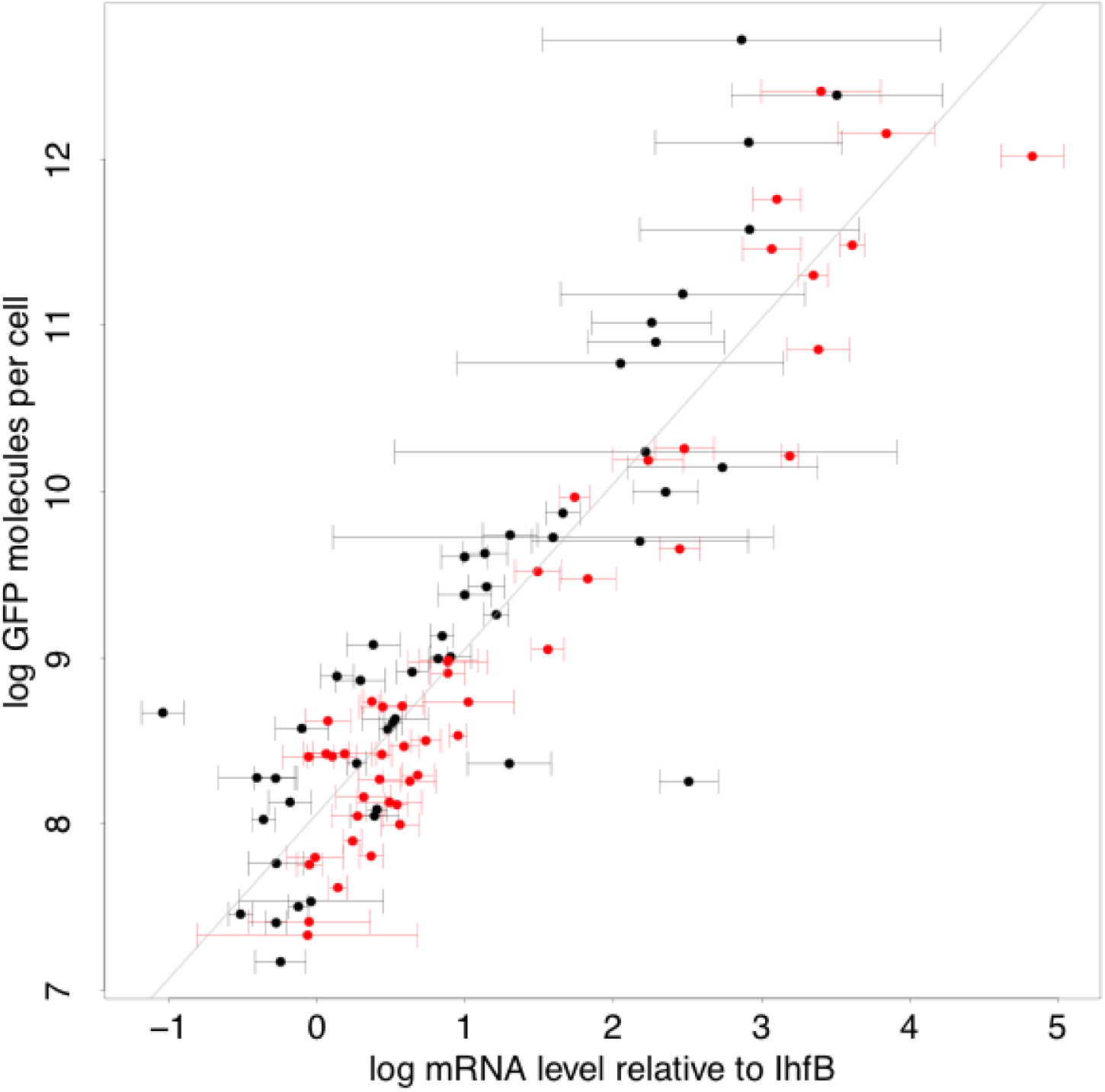
Relationship between log protein levels as measured by GFP intensity in FACS (vertical axis) and log mRNA levels (horizontal axis). The mRNA levels are estimated relative to the mRNA level of reference gene IhfB. Error bars show plus and minus one standard-deviation of the posterior probability distribution on mRNA levels (**Supplementary Text**). Black data points correspond to native promoters and red data points to synthetic promoter. The straight line shows a linear fit with slope 1, i.e. the best fit to a model where the protein level *p* is directly propotironal mRNA level *m*, log(*p*) = *c* + log(*m*), with *c* = 7.06 (**Supplementary Text**).

**Figure S4.**
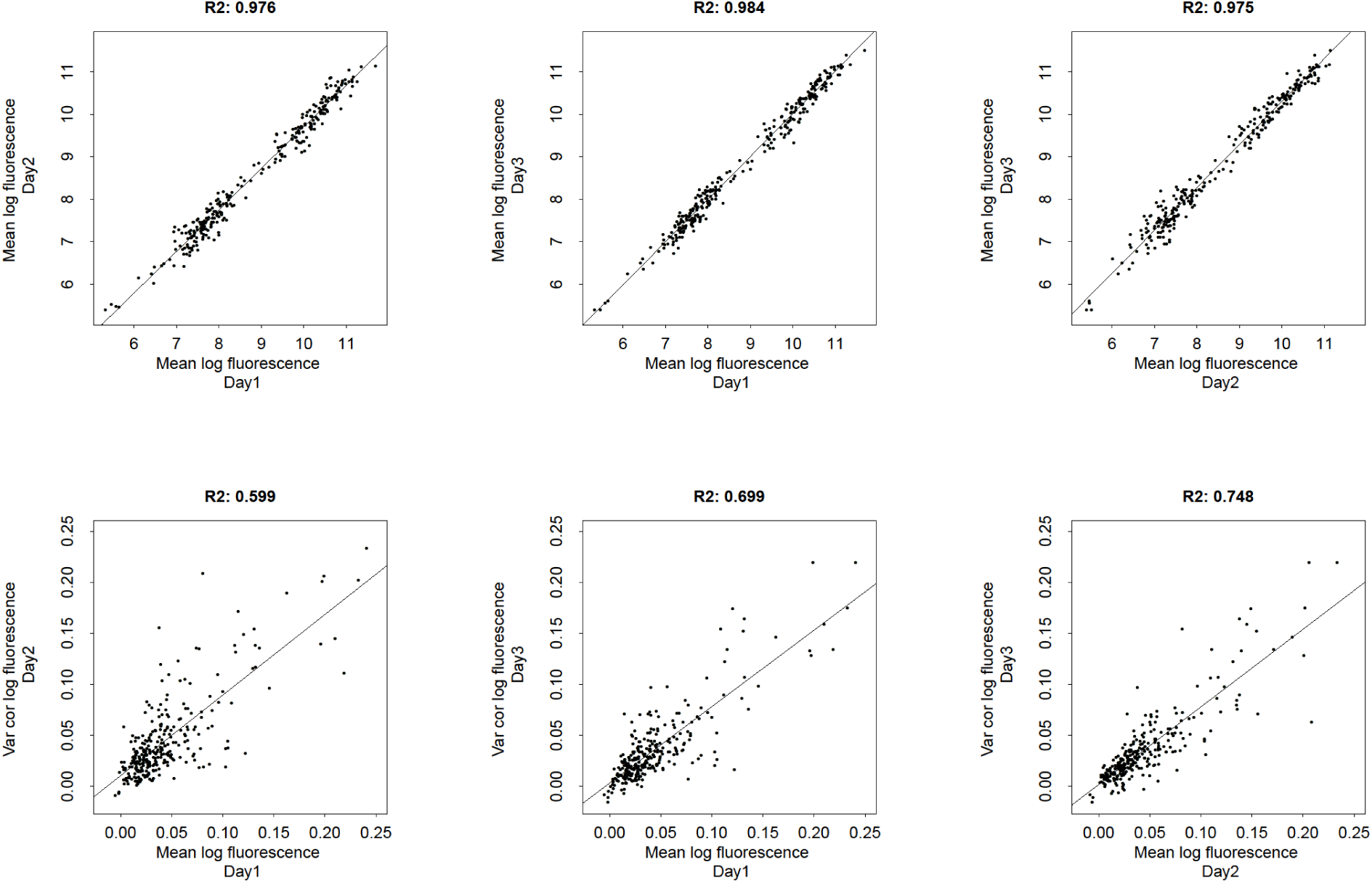
Comparison of three biologial replicate FACS measurements of means and variances of log-fluoresence for evolved *E. coli* promoters. The top 3 panels compare mean log-fluorescences across 3 replicates and the bottom 3 panels compare variances in log-fluorescences across 3 replicates. The Pearson squared-correlation coefficients between pairs of replicate measurements are indicated at the top of each panel.

**Figure S5.**
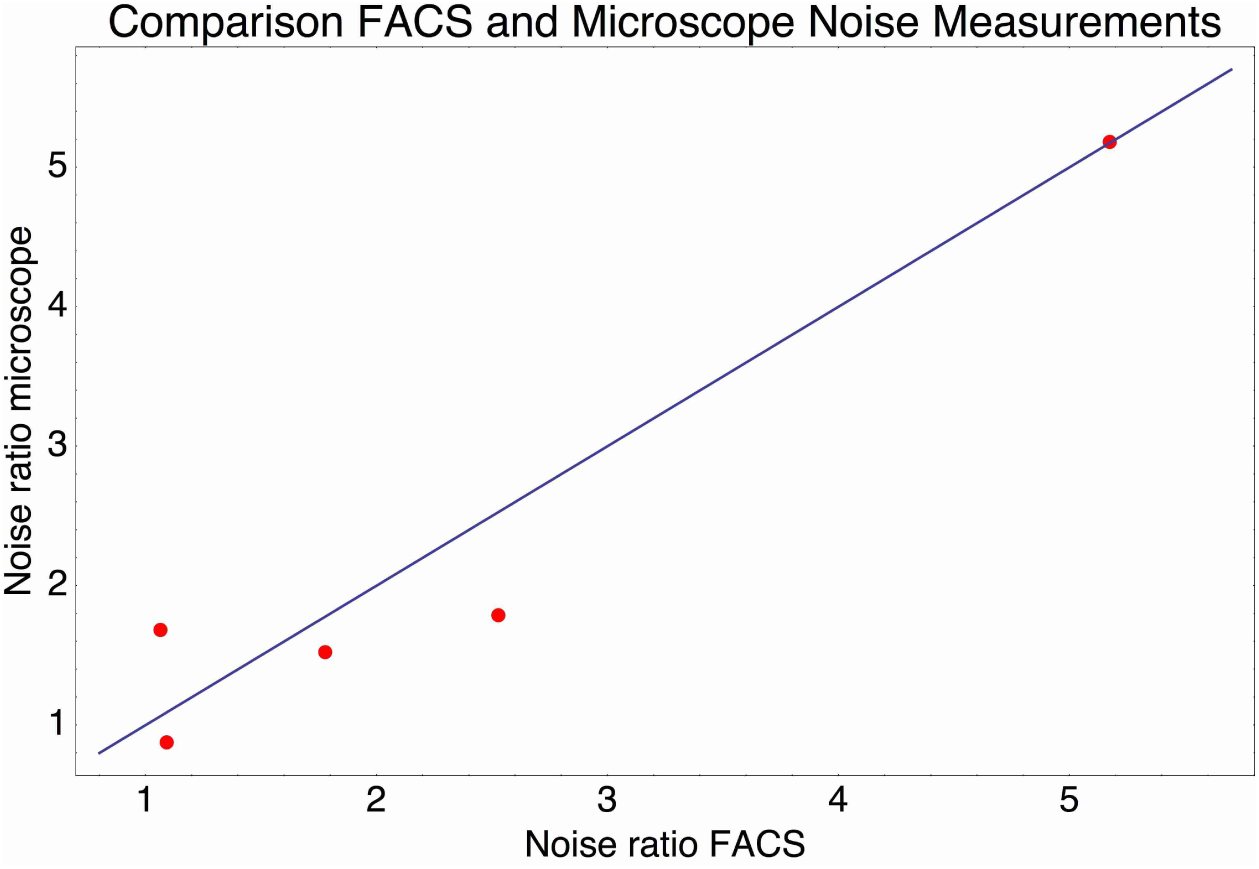
Relative noise levels (variance of the log-expression distribution) of 5 pairs of native promoters that have very similar mean expression levels. Each dot correspond to one of the pairs of promoters and shows the ratio of the noise level of the highest noise promoter to that of the lower noise promoter as measured by FACS (horizontal axis) and by microscope (vertical axis). The blue line shows the line *y* = *x*.

**Figure S6.**
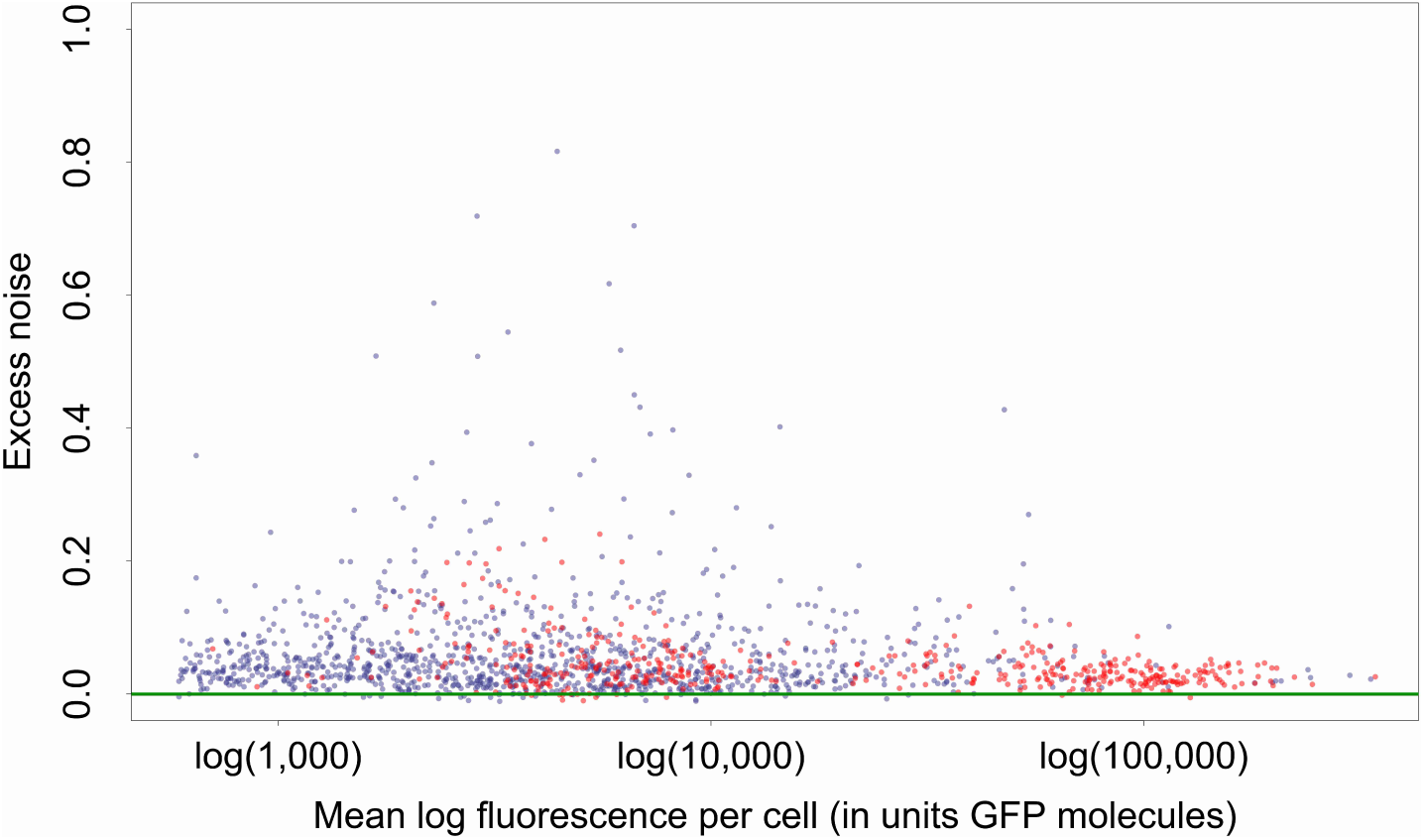
Mean log-fluorescence (horizontal axis) and excess noise levels (vertical axis), i.e. the different between variance of log-fluorescence levels and the minimal variance at the corresponding mean, for all native (blue dots) and synthetic (red dots) promoters. Both axes are in units of number of GFP molecules. Note that, in contrast to raw variances in log-fluorescence, that show a clear dependence on mean log-fluorescence, the excess noise levels show no dependence on mean.

**Figure S7.**
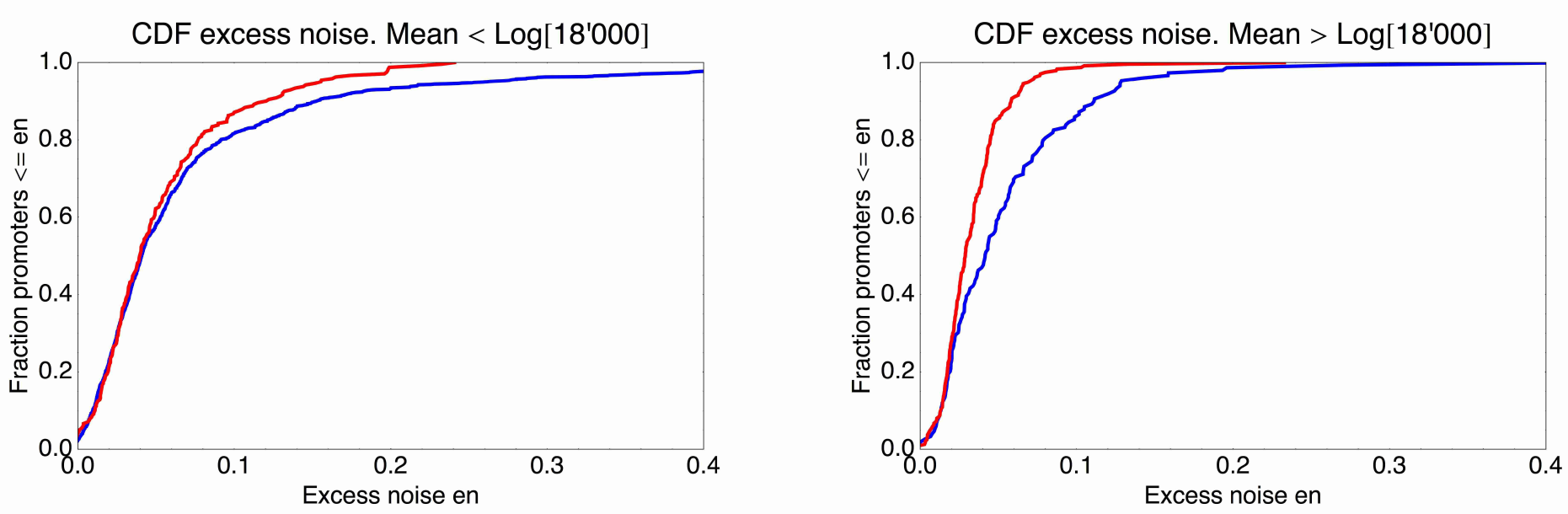
Cumulative distributions of excess noise levels for the native (blue) and synthetic promoters (red). The left panel shows the cumulative distribution of excess noise for promoters whose mean log-expression was less than log(18000) (corresponding to the medium expressing synthetic promoters), and the right panel for promoters with mean log-expression more than log(18000) (corresponding to the high expressing synthetic promoters). High noise promoters are clearly enriched among native promoters for both medium and high expressing promoters.

**Figure S8.**
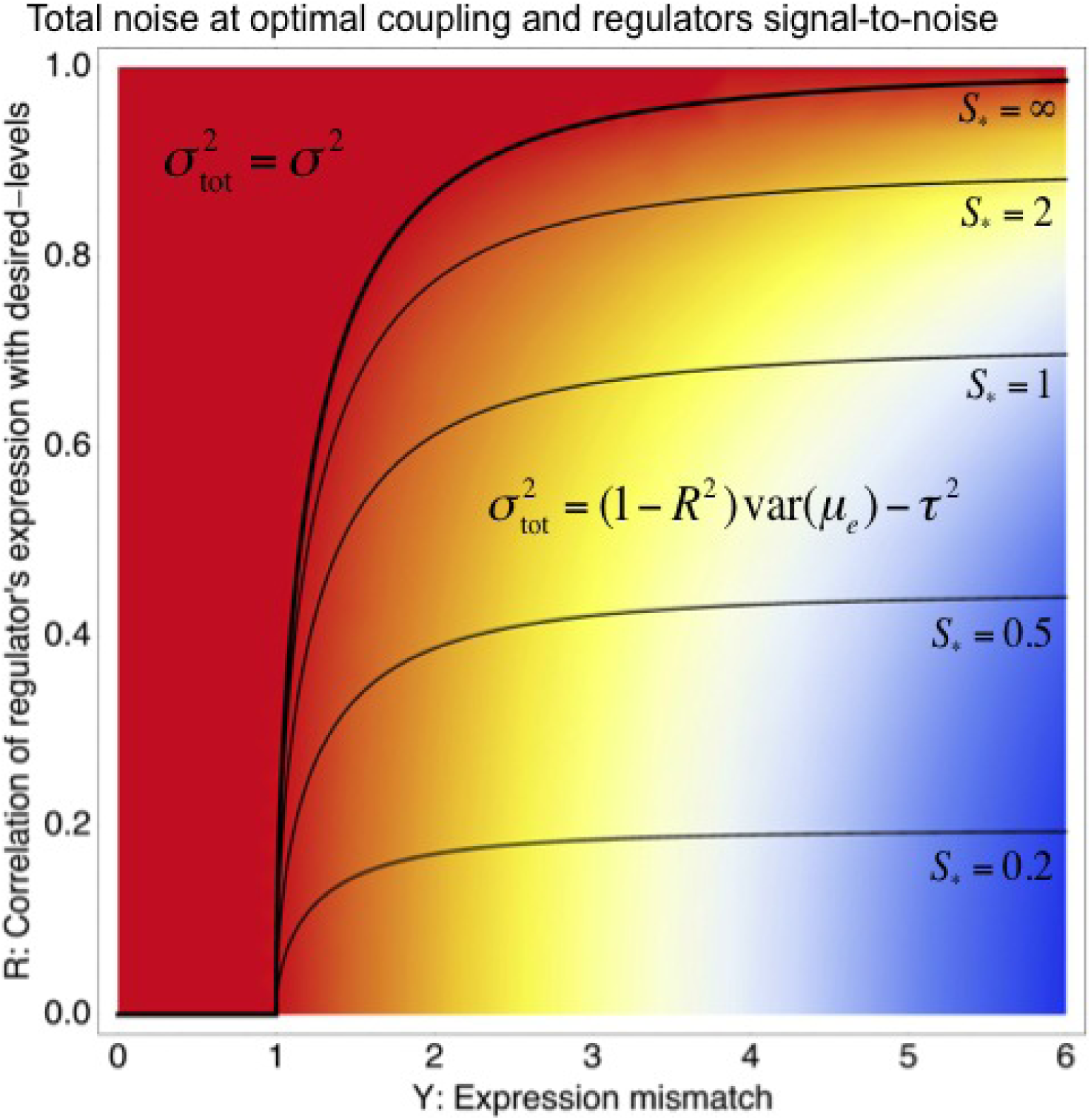
Phase diagram of the total noise *σ*_tot_ of a promoter with expression mismatch *Y* (horizontal axis) that is coupled (at optimal coupling strength) to a regulator whose regulatory activities have correlation *R* with the desired expression levels (vertical axis) and whose signal-to-noise ratio *S* has also been optimized. The colors indicate the value of *σ*_tot_, running from *σ*_tot_ equal to the noise *σ* of the *unregulated* promoter (red) to *σ*_tot_ = 6*σ*. A phase boundary (thick black curve) separates solutions in a ‘basal noise regime’ at the top left, where the total noise equals the minimal noise *σ*^2^, and solutions in an ‘environment-driven noise regime’ at the bottom right, where the total noise matches the variance in desired levels that is not tracked by the regulation, i.e. 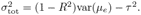 The contours show optimal signal-to-noise ratios *S_∗_* as a function of Y and R. Note that *S_∗_* diverges at the phase boundary.

## Supplementary Text

### Estimating the mean and variance of log-fluorescence levels from FACS data

Visual inspection of the distributions of fluorescence intensities for individual cells containing the same promoter construct shows that almost all of these distributions can be well approximated by a log-normal distribution. We thus chose to characterize the distribution of expression levels of each promoter by the mean and variance of log-fluorescence intensities across cells. Visual inspection of the distributions also indicated that, for almost all promoters, there is a small number of measurements with aberrantly high or low values that are likely due to some measurement artefact, and we designed a Bayesian procedure for automatically discounting these aberrant measurements.

For each clone we typically have around *N* = 5000 independent FACS intensities measured. We denote by *x* the log-intensity (using natural logs) of an individual cell. We first calculate the mean and variance without taking outliers into account, i.e.

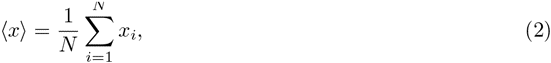

and

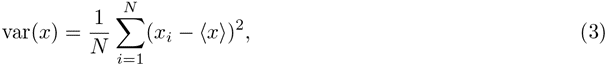

where *x_i_*is the log-intensity of cell *i*. We call these the ‘original’ mean and variance.

Next we take outliers into account. We assume that, of all *N* measurements, only a fraction *ρ* are ‘correct’ measurements, and the other (1 − *ρ*) are ‘outliers’, meaning that these are erroneous measurements. We assume that these ‘outliers’ derive from a uniform distribution that spans the range of measurements *R* = (*x*_max_ − *x*_min_). Finally, we assume that the distribution of ‘correct’ measurements is approximately Gaussian with (unknown) mean *µ* and variance *σ*^2^. Under these assumptions, the probability of a measurement of log-intensity *x* is given by

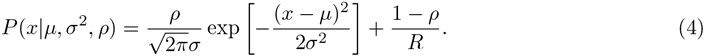

The probability of the entire data-set for a clone is then simply given by

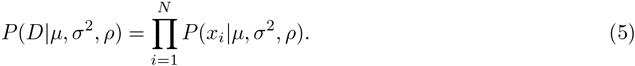

We then maximize this probability with respect to *µ*, *σ*^2^, and *ρ*. This can be easily done using Expectation-Maximization. The resulting mean *µ*, and variance *σ*^2^ are corrected for outliers.

### Inferring the relation between FACS intensity and GFP molecules per cell

To infer the relationship between FACS intensity per cell and GFP molecules per cell we used quantitative Westerns. For each of 8 strains of known FACS intensities, we extracted the protein contents from a fixed number of cells and quantified total GFP intensity. In the same experiment the GFP intensities were measured for known amounts of GFP ranging from 10 to 100 nanograms. We performed 3 replicate experiments. In each replicate we measured the GFP intensity of the 8 strains, as well as ‘reference’ intensities of bands loaded with 10, 25, 50, 75, and 100 nanograms of GFP. We measured intensities from these gels using both 10 second and 20 second exposure times, giving a total of 6 replicate measurements of the reference amounts and the 8 strains.

**Figure S9** shows the measured GFP intensities *I* as a function of the amount of GFP *w* (weight in grams) for the reference bands, in each of the 6 replicate experiments. Note that there are 5 points, corresponding to weights of 10, 25, 50, 75, and 100 nanograms in each curve.

The curves show that the measured intensities are saturating as the amount of GFP increases. Second, the intensity scale varies significantly from replicate to replicate. The simplest linear relationship between *I* and *w* that includes saturation is of the form

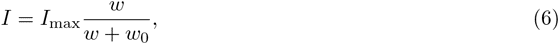

and inspection of the curves shows that each of them can be reasonably well fitted to this functional form. To infer the amount of GFP corresponding for a particular strain in a particular replicate, we need to infer *w* as a function of the measured value *I*. We thus invert the relationship and find the general form

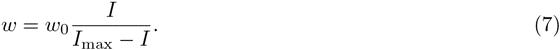

In other words, our functional form assumes that for a suitably chosen value *I*_max_, the weight *w* becomes directly proportional to the transformed variable *I/*(*I*_max_ − *I*). As an example, **Figure S10** shows that for the first replicate, when plotting *w* as a function of *I/*(15631 − *I*), i.e. with a value of *I*_max_ = 15′631, we obtain an approximately linear relationship. Similar approximately linear relationships are observed for the other replicates as well.

**Figure S9.**
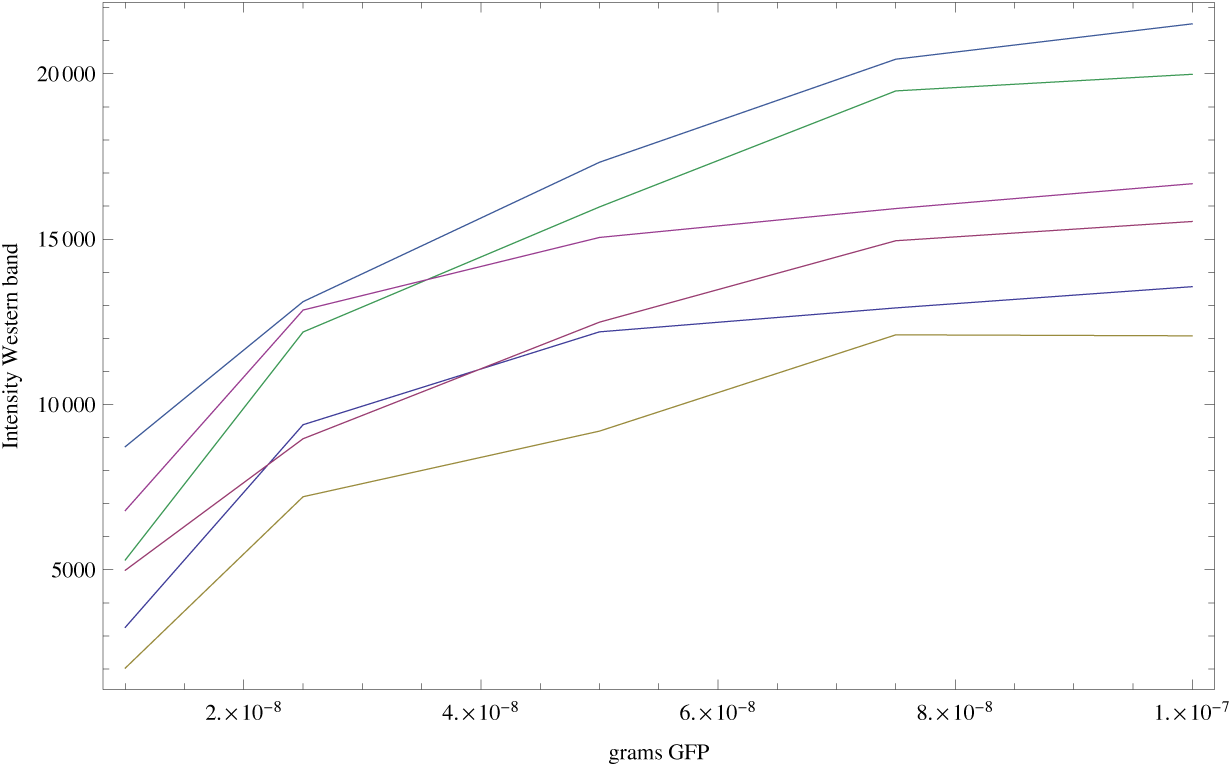
Measured intensities of the GFP reference bands as a function of the amount of GFP (in grams) loaded on each band. Each curve corresponds to one replicate (shown in a separate color), and each curve has 5 data-points.

**Figure S10.**
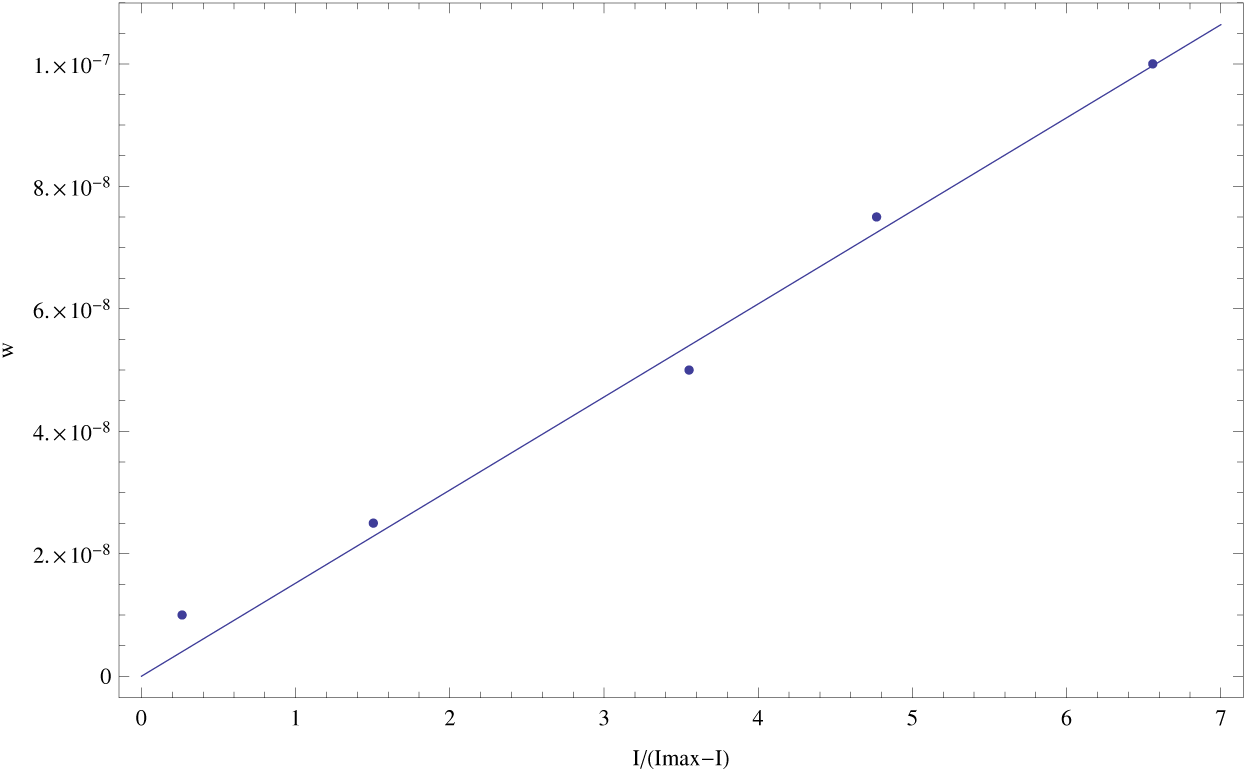
For the first replicate, we inferred a saturation value *I*_max_= 15′631. Plotting w as a function of *I*/(I_max_ − *I*) we obtain an approximately linear relationship that also approximately goes through the origin (0, 0) (as it should).

To fit *w* as a function of *I* for each replicate, we assume that the difference between *w* and *I/*(*I*_max_ − *I*) is Gaussian distributed with unknown variance *σ*^2^. That is, for each data-point *i* in a replicate, the weight *w_i_* and its intensity *I_i_* are related through

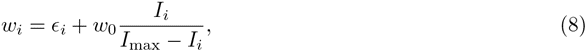

with *∈_i_* the noise, which is Gaussian distributed with unknown variance *σ*^2^, i.e.

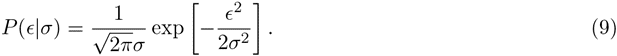

Using this, the probability of the observed data in a titration curve, given parameters *I*_max_, *w*_0_, and *σ* is:

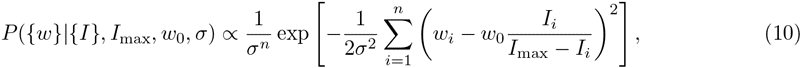

where *n* = 5 is the number of points in a titration curve, and we have ignoted factors of 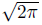 for convenience.

Imagine that we augment our data-set (*{w}, {I}*) with a single data-point (*w_s_, I_s_*), where *I_s_* is the measured intensity of strain *s* and *w_s_*is a hypothesized amount of GFP for this strain. The probability of this entire data-set is given by

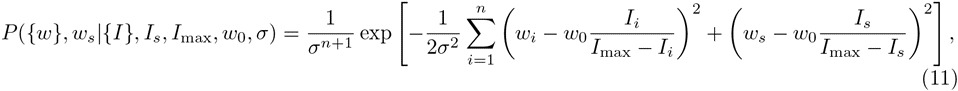

Formally, we now need to specify a prior *P* (*I*_max_*, w*_0_*, σ*) and integrate over these unknown parameters. We will use a uniform prior over *I*_max_ and *w*_0_, and a scale prior 1*/σ* for *σ*. That is, formally we want to calculate

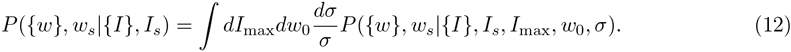

Note, if we additionally integrate over *w_s_* we obtain

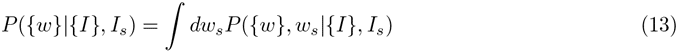

and dividing by this we obtain the posterior distribution of *w_s_*:

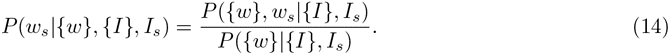

To perform the integrals in (12), we first simplify the notation by denoting the new data-point (*w_s_, I_s_*) as (*w_n_*_+1_*, I_n_*_+1_), i.e. as if it was the (*n* + 1)st data-point. The integrand now takes the form

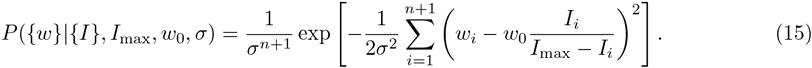

To further simplify notation, we write *y_i_* = *I_i_/*(*I*_max_ *− I_i_*), keeping in mind that the values of *y* depend on *I*_max_. Further, for any quantity *x* that takes on values *x_i_*over the 5 titration points and the added point, we write averages like

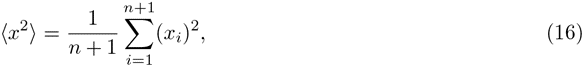

and so on. The integrand can then be rewritten as

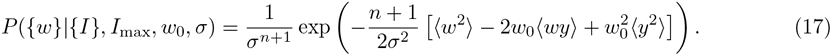

Performing the integral over *w*_0_ we obtain

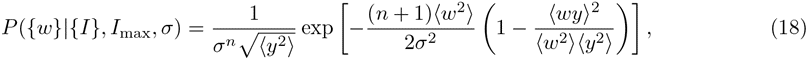

where we have again ignored prefactors that cancel in the final posterior for *w_s_*, i.e. equation (14). We next integrate over *σ*. Performing this integral we obtain

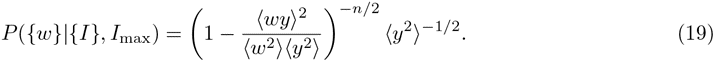

Notice that the key expression in parentheses is simply one minus the squared correlation coefficient between the variables *w* and *y*, i.e.

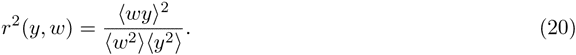

In other words, the values of *w_s_* and *I*_max_ that maximize the probability are those such that the linear correlation between the resulting values of *y* and the *w* values is maximal.

We next need to perform the integral over *I*_max_. Since this integral cannot be performed analytically, we performed the integrals over *I*_max_ numerically, separately for each strain *s* and each of the 6 replicates. Finally, for each replicate *r* and strain *s*, we determined the values *w_rs_* that has maximal posterior probability. These are our estimated GFP amounts (in grams) for each strain and replicate (**Fig. S11**).

Although the inference clearly separates the high expressed from the low expressed clones, curves from the different replicates seem to be separated by constant shifts from each other. Since the vertical axis is shown on a logarithmic scale, this means that the curves differ by common multiplicative factors. This difference in scale is almost certainly due to an experimental artefact and we will thus normalize for it.

**Figure S11.**
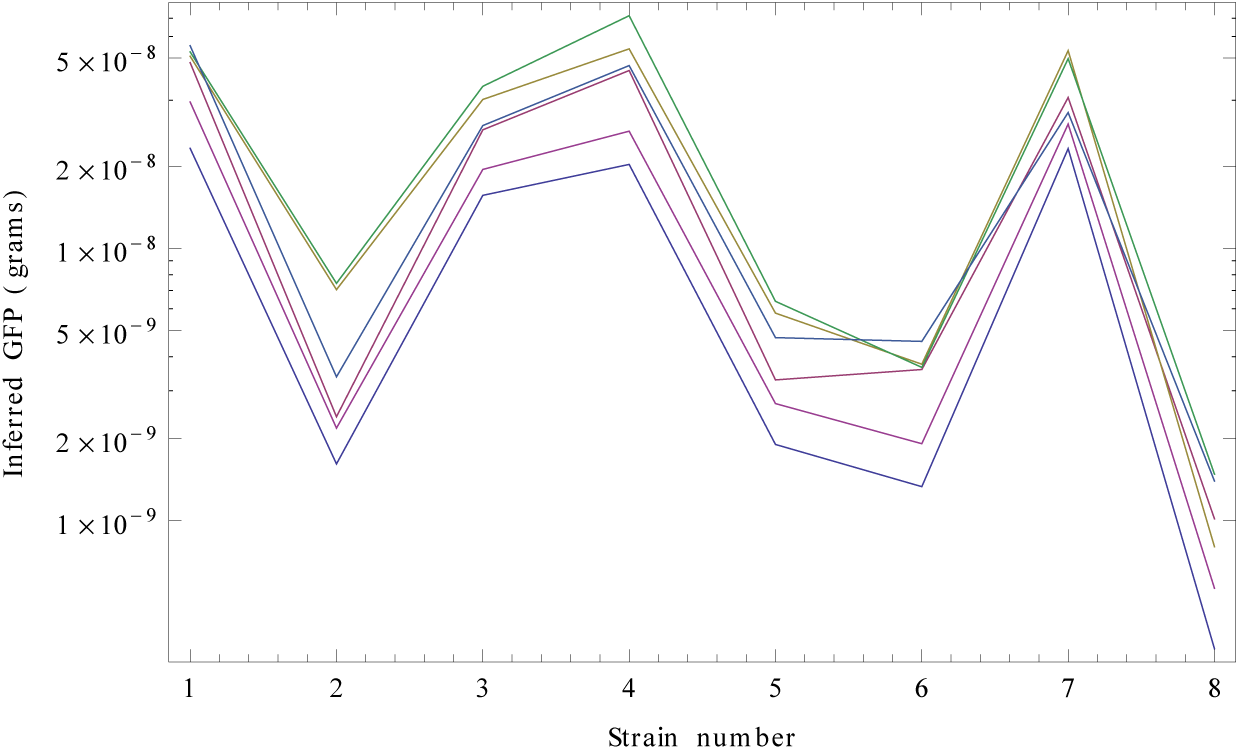
Inferred GFP amounts (in grams, vertical axis) for the 8 strains (strain numbers shown along the horizontal axis) using the reference data from each replicate. Each color corresponds to a replicate. The vertical axis is shown on a logarithmic scale.

Let *w_s_*(*i*) be the inferred amount of strain *s* in replicate *i*. To account for the variability in overall scale, we normalize the inferred log-weights in each replicate by calculating the average log-weight in the replicate, i.e.

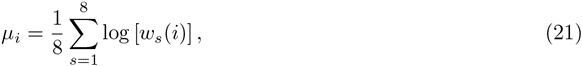

and a total average scale of the replicates

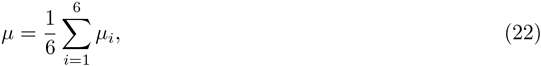

and then transforming the estimated shifts as follows:

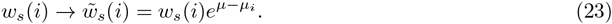

In addition, dividing the weight 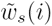 by the known weight of a single GFP molecule (4.48210*^-^*^20^ grams), we get an estimate of the number of GFP molecules in the bands for each strain. Finally, we used OD measurements to estimate the number of cells loaded on each band, and divided by these to obtain an estimate of the number of GFP molecules per cell for each of the strains across each of the replicates. **Figure S12** shows the inferred GFP molecules per cell for each strain after normalization, which indeed show much less variation across the replicates.

**Figure S12.**
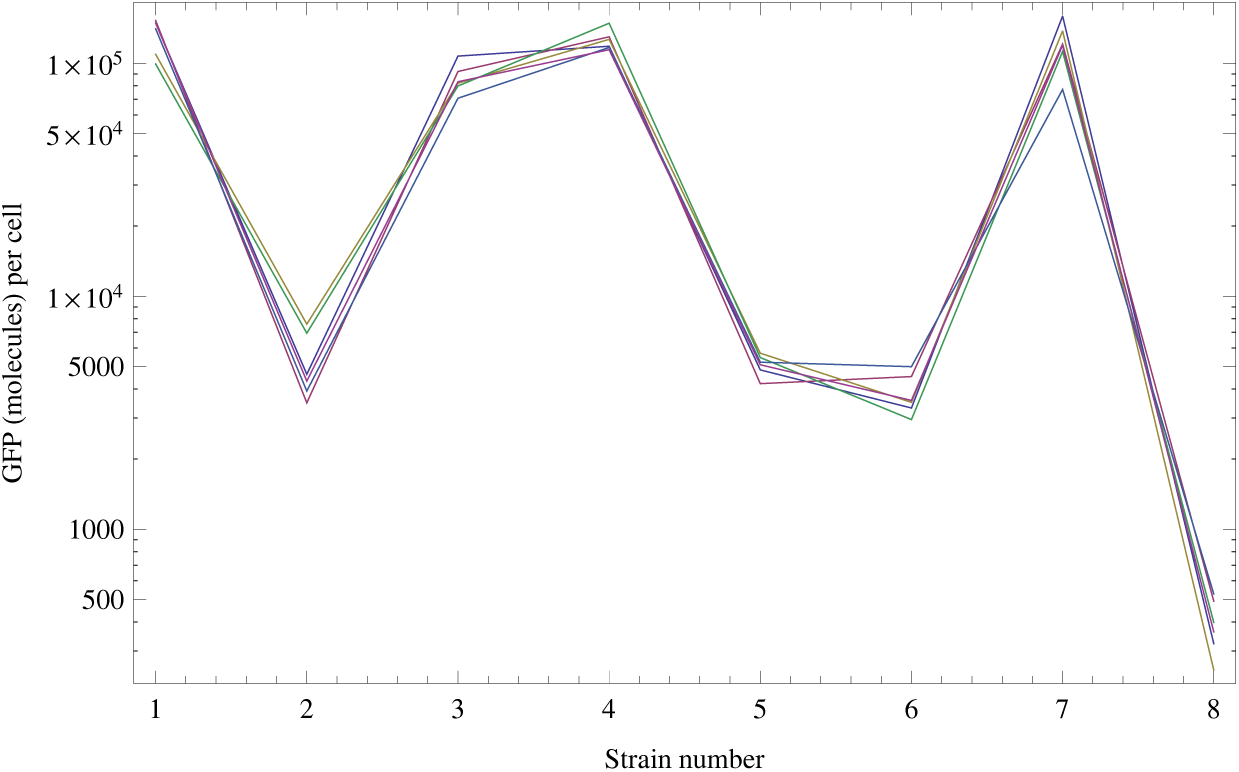
Normalized inferred GFP amounts (molecules per cell, vertical axis) for the 8 strains (strain numbers shown along the horizontal axis) using the reference data from each replicate. Each color corresponds to a replicate. The vertical axis is shown on a logarithmic scale.

Finally, we compare the inferred GFP amounts for each strain, with the FACS intensities measured for that strain. Observing that the variation in both estimated FACS intensities and GFP molecules per cell increases with the mean, it is most natural to compare GFP and FACS levels on a logarithmic scale. Let *f_s_* denote the true log-FACS intensity and *g_s_* denote the true log-GFP molecules per cell. Assuming that GFP molecules per cell and (background corrected) FACS intensity are directly proportional to each other, the log-levels are related through

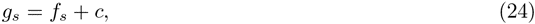

with *c* a constant. For each strain *s*, we calculated the mean log-FACS intensity 〈*f_s_*〉 and its variance var(*f_s_*) across replicate FACS, as well as the mean log-GFP molecules per cell 〈*g_s_*〉 and its variance var(*g_s_*) across the quantitative Westerns as described above. Assuming Gaussian deviations between the true and observed levels, the probability of the data given *c* is given by

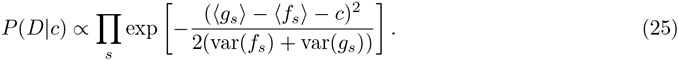

We thus find for the optimal value of *c*:

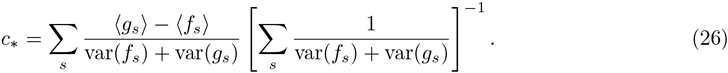

**Suppl. Figure 2** shows the estimated log-FACS and log-GFP levels including their error bars, together with the optimal fit *c_∗_* = 1.06.

Consequently, if *F* is the FACS intensity of a strain (non-log), then the estimated number of GFP molecules per cell *G* is equal to *G* = *e*^1^*^.^*^06^*F* = 2.88*F* . Note that, with these estimates, the highest expressed strain, with an average FACS intensity of 37′500, would have about 108′000 molecules of GFP per cell. The lowest expressed strain (with FACS intensity 143) would have 415 molecules per cell. From now on, we will multiply all FACS intensities by 2.88 so that a FACS intensity of *I* automatically corresponds to the fluorescence of *I* GFP molecules, i.e. we express FACS fluorescence intensities in units of GFP molecules per cell.

### Comparing mRNA and protein levels

For 94 clones, we quantified mRNA levels using qPCR. The qPCR procedure uses a standard reference curve which allows it to infer the absolute number of molecules of the mRNA of interest in the input sample. Each input sample is created by extracting RNA from a certain number of cells (which we can estimate approximately), and reverse transcribing this RNA into cDNA. Unfortunately, both the total amount of cells used, as well as the efficiency of the reverse transcription can fluctuate significantly outside of our control, and this will make the total number of molecules detected fluctuate as well. To control for this, we always quantify the absolute number of molecules of two types of mRNAs in parallel for each sample; the mRNAs of the gene of interest, and the mRNAs of a reference gene which we are confident is constantly expressed. The reference gene we used was *ihfB*.

For each promoter of interest *p*, we obtained measured mRNA molecule numbers together with mRNA molecule numbers for the reference gene, in 3 separate biological replicates, and in 3 technical replicates for each biological replicate, i.e. 9 pairs of measurements in total. We denote the log-quantity of the mRNA of promoter *p* in biological replicate *r* and technical replicate *i* as *x_pri_*, and the log-quantity of the reference gene in the same replicate as *y_pri_* (note that this depends on the promoter *p* because these quantities come from a common sample). To estimate a single log-ratio *x_p_* − *y_p_* between the expression of the gene of interest and the reference gene, we will proceed as follows. First, we will integrate the data from the technical replicates to obtain biological replicate expression *x_pr_* and *y_pr_*. We then combine the differences *d_pr_* = *x_pr_* − *y_pr_*across the biological replicates, to obtain the final *d_p_* = *x_p_ − y_p_*.

The statistical model that we use assumes that the difference between the value *x_pri_* measured in technical replicate *i*, and the true expression *x_pr_* is Gaussian distributed with mean zero and an unknown variance 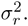 Note that we assume that this ‘noise’ is the same for all promoters *p*, but may fluctuate between biological replicates. We similarly assume the difference between *y_pri_* and *y_pr_*is Gaussian distributed with variance 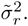 We noted that there is a small fraction of measurements that deviate by large amounts from the measurements in other replicates. We assume that there is a small fraction of measurements that failed in some way, giving erroneous measurement values. To take this into account we will use a mixture model that asssumes a small fraction of the measurements come from a uniform distribution that spans the observed range of the data.

Let *R_r_* = max*_p,i_*(*x_pri_*) − min*_p,i_*(*x_pri_*) denote the range of observed values in biological replicate *r*, and let *ρ_r_* denote the fraction of measurements in replicate *r* that are meaningful, i.e. not erroneous. The probability of a single measurement *x_pri_* given *x_pr_*, the variance 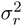 and fraction *ρ_r_*is given by

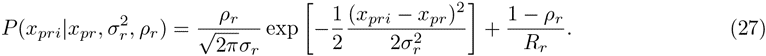

The probability of all technical replicates for all promoters is then simply given by the product over all promoters *p* and technical replicates *i*:

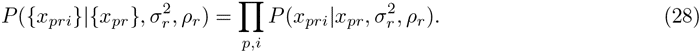

We next maximize this likelihood with respect to the fraction *ρ_r_*, the variance 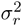 and the expression levels *x_pr_* for all promoters *p*. This optimization can be done using a straight-forward Expectation Maximization scheme.

### Expectation Maximization

Given a current estimate of *x_pr_*, of the variance 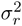 and the fraction *ρ_r_*, the posterior probability that the technical replicate with value *x_pri_* was a meaningful measurement is given by

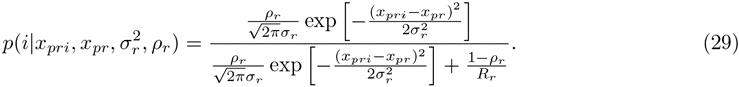

Using these posteriors, the updated value 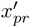 is given by the mean of the technical replicate measurements, weighted by their posteriors

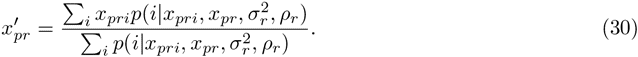

Given current values of *ρ_r_* and 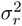 we use these equations to iteratively update all *x_pr_* until they converge. We then update the values of *ρ_r_* and 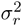 using the following equations

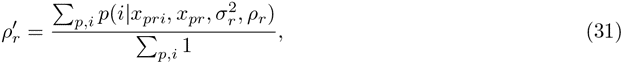

i.e. the updated 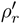 is the average of the current posteriors over all promoters and technical replicates. The update equation for the variance is given by

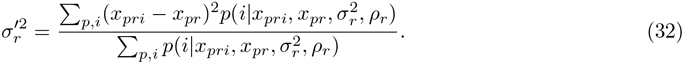

After each update of 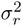 and *ρ_r_* all *x_pr_* are updated until convergence again, and this is iterated until the 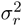 and *ρ_r_* converge. Exactly analogous expectation-maximization equations are used to optimize the values 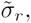 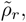 and all *y_pr_* of the reference gene measurements.

Table S1 shows the fitted fractions and variances for each of the replicates. We see that for the majority of replicates the noise level lies around 0.01 (meaning a measurement error-bar of about 0.1 on log-expression), but it is two and three-fold higher for measurements of the genes of interest in two replicates. The fraction of correct measurements ranges from about 93% to almost 100% across the replicates.

**Table S1.**
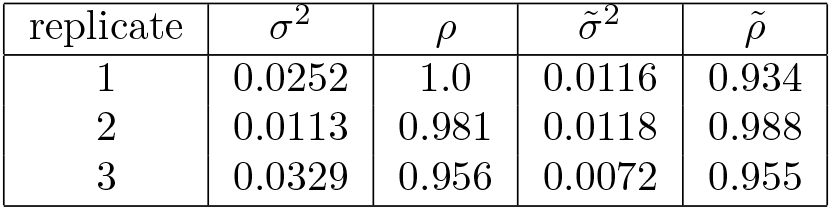
Fitted variances and fractions of meaningful measurements for the genes of interest (σ2, ρ) as well as for the reference gene measurements (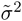, 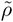) for each of the three biological replicates.

| replicate | $\sigma^2$ | $\rho$ | $\tilde{\sigma}^2$ | $\tilde{\rho}$ |
| --- | --- | --- | --- | --- |
| 1 | 0.0252 | 1.0 | 0.0116 | 0.934 |
| 2 | 0.0113 | 0.981 | 0.0118 | 0.988 |
| 3 | 0.0329 | 0.956 | 0.0072 | 0.955 |

When the variances and fractions have been optimized, we obtain the final technical replicate-averaged quantities *x_pr_*and *y_pr_* and we determine final variances 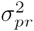 and 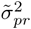 for each of these averages. These final variances are calculated as follows. For each promoter *p* and each biological replicate *r*, we determine the effective number of correct measurements as

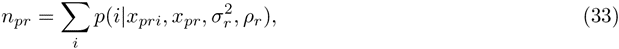

and the final variance is then given by

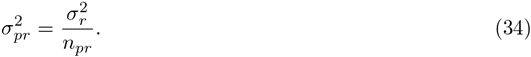

Analogously, for the reference gene measurements we have

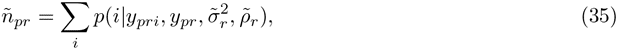

and the final variance

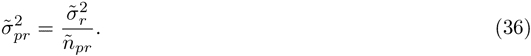

### Combining the biological replicates

For each promoter *p*, we want to estimate the log-expression ratio *x_p_* − *y_p_*by combining the estimated values *x_pr_* and *y_pr_* from each of the replicates, taking into account that these values have different variances for different replicates. For a protein *p* and replicate *r*, the estimated log-expression difference *d_pr_* is

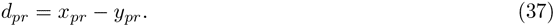

The variance *τ*^2^_*pr*_ associated with that estimated difference is

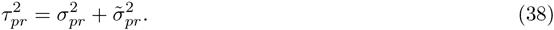

Inspection of the variation in *d_pr_* across biological replicates, relative to their uncertainties *τ_pr_*, makes it clear that, in addition to the uncertainty in each of the estimates *d_pr_*, there is substantial variation in *d_pr_* across the biological replicates which is quite different for different promoters. That is, for some promoters the biological replicates give very consistent *d_pr_*, lying within the error-bars *τ_pr_*, whereas for other promoters the variation in *d_pr_*is much larger than the error-bars *τ_pr_*, indicating that there must be additional variance across replicates.

We will assume that the true value 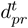 is given by the mean *d_p_*for the promoter plus a biological replicate variation *δ_pr_*

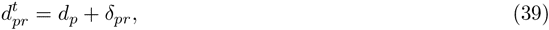

and we will assume that the deviation *δ_pr_* is Gaussian distributed with mean zero and unknown variance 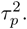 The probability of the estimate *d_pr_*given its variance 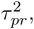 the true value *d_p_*, and the biological replicate variance 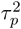 is given by

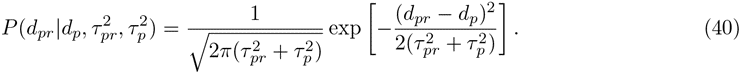

The probability of the data combining all biological replicates for the promoter is simply

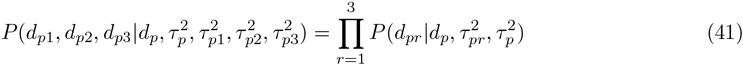

For each promoter *p*, we now maximize this probability with respect to both 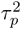 and *d_p_*. Given a fixed value of 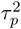, the optimal value of *d_p_* is given by the weighted sum

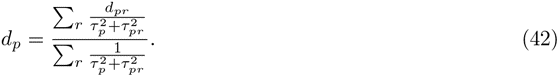

Substituting this optimal *d_p_* into the probability (41), the expression becomes a function of the variance 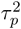 only, and we numerically determine the optimal value of 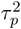 for each *p*. In this way we obtain a final estimate *d_p_* for each promoter. The variance 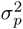 associated with this final estimate is given by

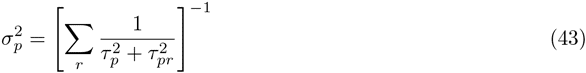

**Suppl. Fig. S3** shows the relationship between protein levels (estimated by FACS) and estimated mRNA levels for the 94 strains for which we measured mRNA levels using qPCR. We see there is a very good correlation between protein and mRNA levels (Pearson correlation *r*^2^ ≈ 0.82). Note that, for a given promoter, the average protein level *p* is related to average mRNA level *m* by the ratio of the translation rate *λ* and protein decay rate *µ*. That is,

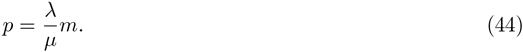

Since GFP is very stable compared to the duplication rate of our cells, for our system the protein decay rate *µ* is approximately equal to the growth rate of the cells, and thus constant across the promoters. Consequently, the fact that 82% of the variation in protein levels is explained by variations in mRNA levels, suggest that the translation rate *λ* shows relatively small variations across the strains. Below, we use this data to more rigorously estimate variation in translation rates across the strains.

### Estimating relative translation rates

As before, we denote by *d_p_* the relative (to *ihfB)* log-mRNA level of promoter *p*, and we will denote by *y_p_* the log protein number per cell (as measured by FACS) for promoter *p*. Denoting by *m* the absolute number of mRNAs per cell for the reference gene *ihfB*, by *λ* the average translation rate, and by *µ* the protein decay rate (as a consequence of cell growth), *y_p_* and *d_p_* are related through

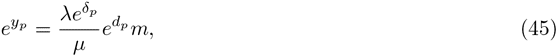

where we have written the translation rate *λ_p_*of promoter *p* in terms of the average translation rate *λ*, and a promoter specific deviation *δ_p_*, i.e. 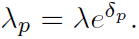 Defining *e^c^*= *λm/µ* we have

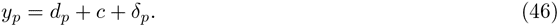

Using that our estimate of *d_p_* is Gaussian distributed with standard-deviation *σ_p_*, and assuming that *δ_p_*is Gaussian distributed with mean 0 and standard-deviation *τ*, the probability of our data given the *σ_p_*, *c*, and *τ* is

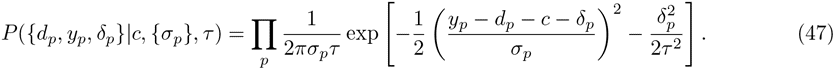

To estimate the variance *τ*^2^ we integrate over all *δ_p_* and *c* (using a uniform prior). To simplify the notation of the result we write 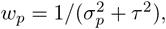 and Δ*_p_* = *y_p_* − *d_p_*. We then have

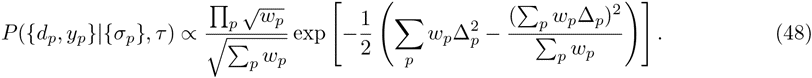

We numerically determine the value of *τ* that maximizes this likelihood and find *τ_∗_* = 0.47. Using this maximum likelihood value of *τ*, the maximum likelihood value of *c* is given by

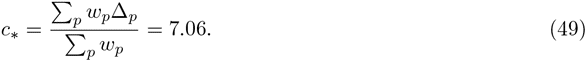

The fit *y* = *c* + *d* is shown as the black line in **Suppl. Fig. S3**.

Finally, using *τ_∗_* and *c_∗_*, we determine the most likely values of the *δ_p_*. We find

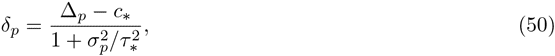

with a standard-deviation of

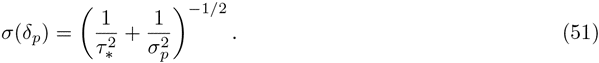

**Figure S13** shows the estimated values of *δ_p_*, together with their error bars *σ*(*δ_p_*), as a function of the log protein level *y_p_*. We see that, for the large majority of promoters, the estimated translation rate is within 2 − 3 fold of the average translation rate (i.e. |*d_p_*| < 1), confirming that there is relatively little variation in translation rates. For the most extreme example, the translation rate is approximate *e*^1^*^.^*^9^ = 6.6 fold lower than the average translation rate.

**Figure S13.**
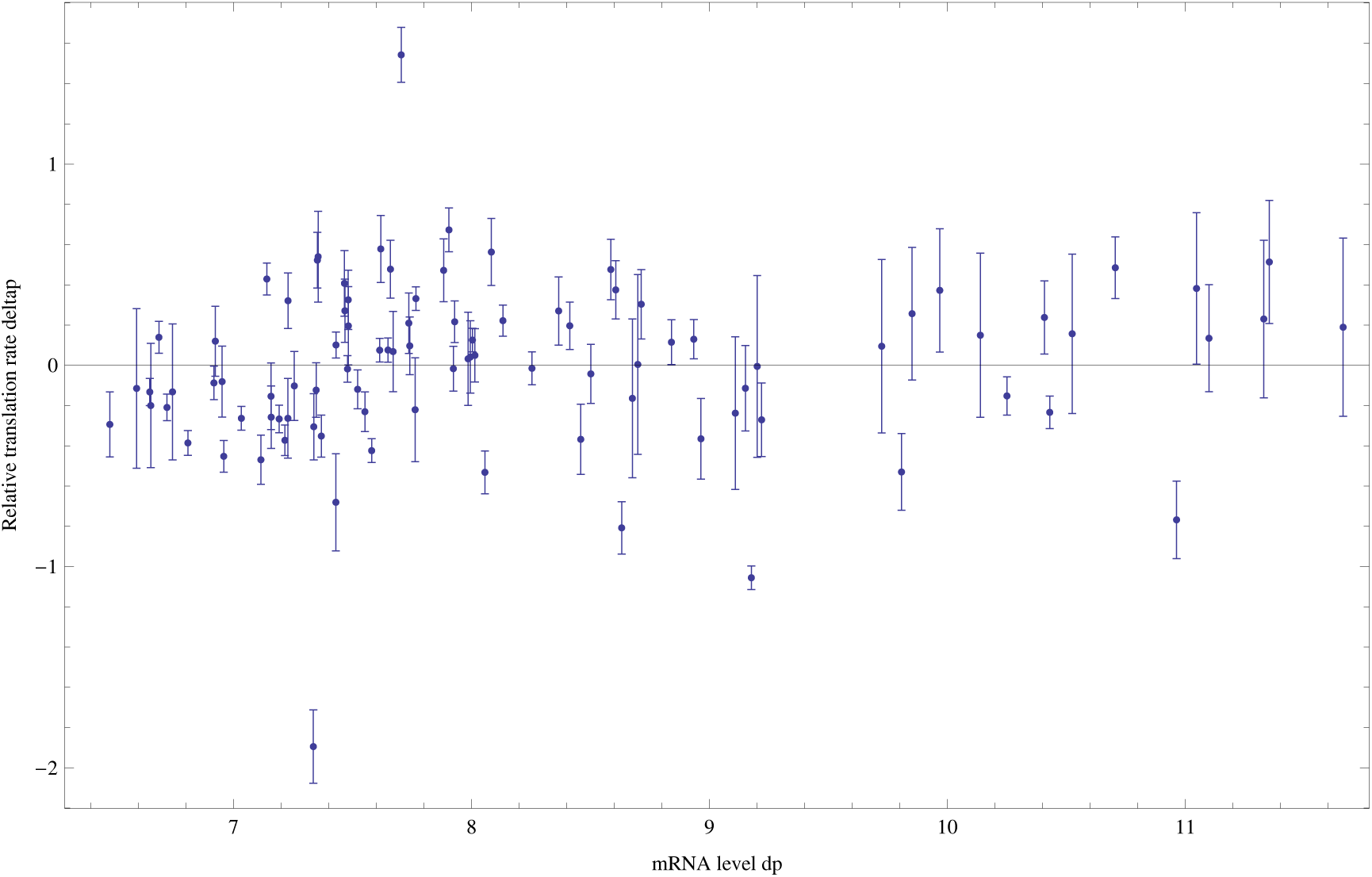
Estimated relative log-translation rates *δ_p_* and their error bars *σ*(*δ_p_*) (vertical axis) as a function of the log-mRNA level relative to *ihfB*, *dP*, for each promoter *p*.

The figure also shows that there is no correlation between the relative translation rate *δ_p_* and the log mRNA level *d_p_*. We also find no correlation of *δ_p_*with either log protein level *y_p_*, or the variance of the log protein level (data not shown).

### Minimal expression noise as a function of mean expression

To model the noise distribution of our promoters we start with the simple case in which there are constant rates of transcription, translation, mRNA decay, and protein decay. Let *λ_m_* be the rate of transcription per unit time, µ*_m_*the rate of mRNA decay (per mRNA per unit time), *λ_p_* the rate of translation (per mRNA per unit time), and *µ_p_* the rate of protein decay (per protein per unit time). Note that in our case all proteins decay at the same rate and, because the decay rate of GFP is relatively small compared to the dilution rate as a consequence of cell growth, the rate *µ_p_* is effectively given by the growth rate of the cells.

In [25] an analytical expression was derived for the distribution *P* (*n*|*λ_m_*, µ*_m_*, *λ_p_*, *µ_p_*) under the assumption that the rate of protein decay is *small* compared the rate of mRNA decay. In E. coli the typical mRNA decay rate is on the order of 5 minutes [26]. In the minimal media with glucose in which our cells are grown, the doubling time is more than half an hour, so that the protein decay is indeed smaller than the mRNA decay rate by a factor of approximately 6. Since this is not a very large factor, one may worry that for stable mRNAs the approximation breaks down. Fortunately, in [25] it was also shown (by simulation) that as long as the mRNA decay rate is *at least* as large as the protein decay rate, then the approximation still is quite accurate. We will thus assume that we can use this approximation.

Under this approximation the stationary distribution of the number of proteins per cell depends only on the following two ratios:

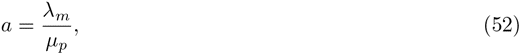

and

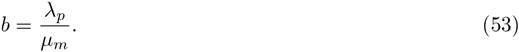

The ratio *a* gives the expected number of transcripts that are produced during the life-time of a single protein, which in our case effectively means the doubling time of the cells, i.e. *a* is the expected number of transcription events per cell cycle. The ratio *b* gives the expected number of proteins that are produced from a single mRNA during its life-time. This is sometimes referred to as the ‘burst size’. That is, typically one assumes *b >* 1 and given that mRNA decay is faster than protein decay, the proteins are produced ‘fast’ from a single mRNA in comparison to the life-time of a typical protein, i.e. in a burst.

The limit distribution *P* (*n|a, b*) is given by a negative binomial

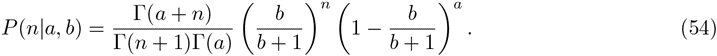

This distribution has a mean

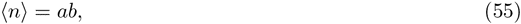

and variance

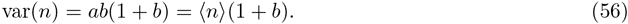

We extend this simple model by assuming the ratios *a* and *b* fluctuate themselves (most likely on a somewhat slower time scale). Although we will not attempt to specify the molecular origins of these fluctuations in *a* and *b*, they likely include fluctuations in the concentrations of polymerases, ribosomes, and transcription factors that regulate the promoter in question. Such fluctuations would contribute to the extrinsic noise of the promoters, since they would equally effect two copies of the same promoter in the same cell. However, they may also include fluctuations in the state of the promoter itself and such fluctuations would contribute to the intrinsic noise.

We will assume that the fluctuations in these ratios of rates are multiplicative, i.e. proportional to the means 〈*a*〉 and 〈*b*〉:

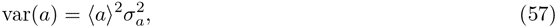

and

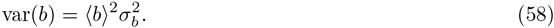

We then find for the total variance of *n*

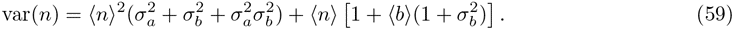

To simplify notation, we introduce the variable

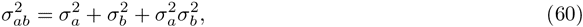

and the renormalized burst-size

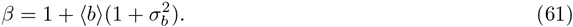

With these definitions we have

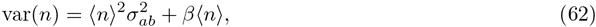

which brings out most clearly that there is a term proportional to 〈*n*〉^2^ that results from fluctuations in *a* and *b*, and a term proportional to 〈*n*〉 that results from Poisson fluctuations in mRNA and protein production, and is proportional to the burst-size.

We now want to relate this expression to variations in log-fluorescence intensities as measured using FACS. Here it is important to note that the log-fluorescence intensity per cell is the result of a combination of fluorescence coming from GFP proteins and *background* fluorescence of the cell.

### Background fluorescence

To estimate background fluorescence, we performed 3 replicate measurements of populations of cells without any plasmid, and 3 replicate measurements of populations of cells containing an empty plasmid (not containing a GFP gene). **Figure S14** shows the reverse cumulative distributions of observed intensities in these control populations (colored lines).

**Figure S14.**
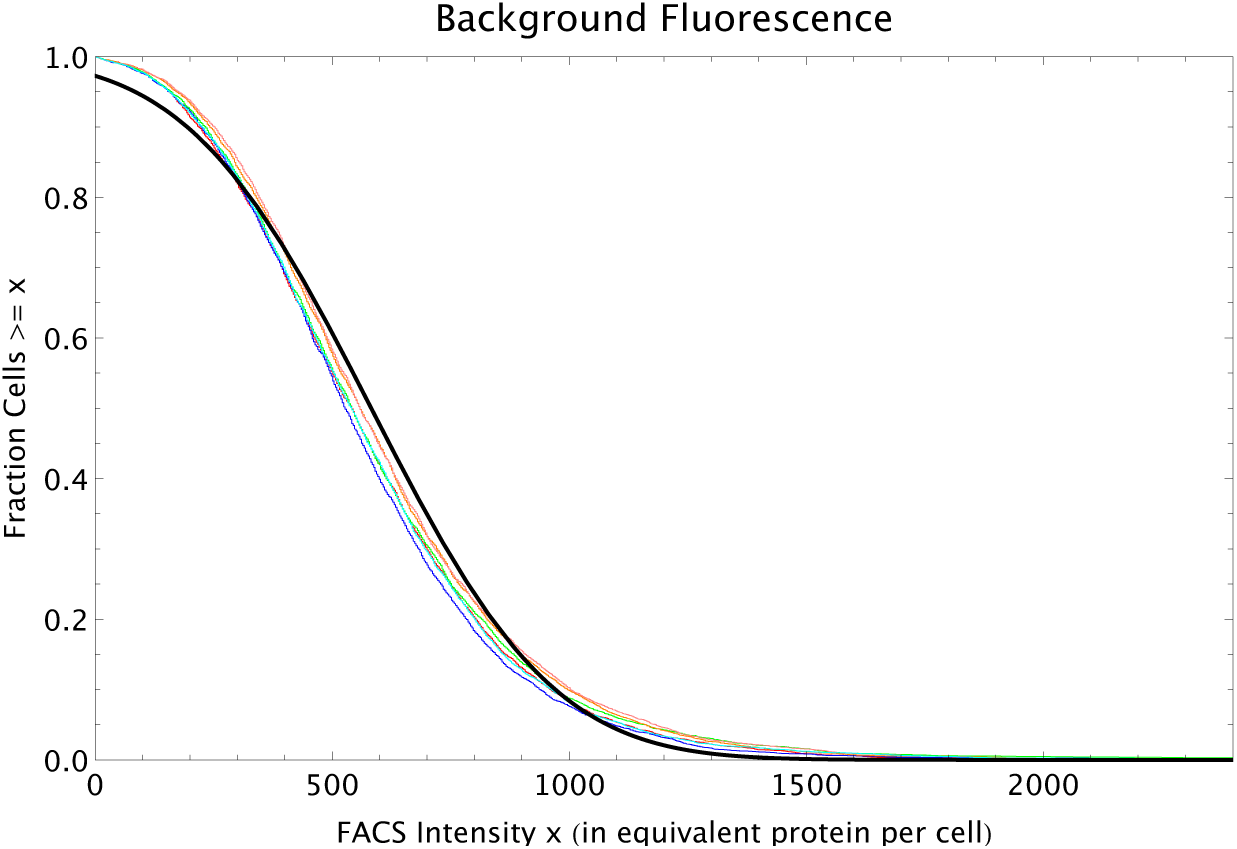
Reverse cumulative distributions of the FACS intensities per cell (multiplied by 2.88 so as to correspond to the equivalent of GFP proteins per cell) for MG1655 cells without a plasmid (red, blue and green curves) and MG1655 cells with an empty plasmid (orange, pink and cyan curves). The black line shows a Gaussian distribution with matching mean and variance.

The curves show that each replicate shows a highly similar distribution of fluorescence levels, and pooling the data from all replicates we find a mean background fluorescence of *n*_bg_ = 582.3 with a standard-deviation of *σ*_bg_ = 302.9. As shown by the black curve in **figure S14**, the distribution of background fluorescences is reasonably approximated by a Gaussian with the same mean and standard-deviation.

### Relating measured variations to the theoretical expression

Let *n*_meas_ denote the measured FACS intensity of a cell. We will write this measured intensity as the sum of an average background fluorescence *n*_bg_, the average number of proteins 〈*n*〉, and a fluctuation of size 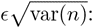

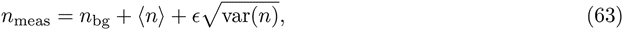

Here *∈* is a quantity that fluctuates from cell to cell, which has mean zero 〈*∈*〉 = 0, and variance one, i.e. 〈*∈*^2^〉 = 1.

We will assume that the fluctuations 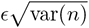 are small re lative to the mean 〈*n*_meas_〉 = *n*_bg_+ 〈*n*〉.

We can then write for the logarithm of the measured FACS intensity

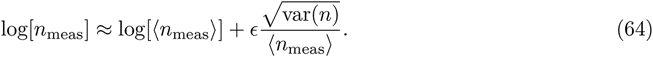

We then find for the variance in log-scale of the measured FACS intensities

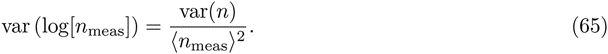

If we substitute the expression (62) for the numerator, we obtain

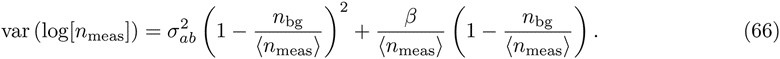

The left panel of **figure S15** shows the mean and variances of the log-FACS intensities of all native promoters. This scatter shows that, as a function of the mean FACS intensity, there is a sharp lower bound on the observed variances. The red curve shows that this lower bound can be well-fitted by a function of the form (66), where we used parameters 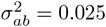 and *β* = 450. Note that the value of 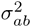 determines the variance in the limit of large means, whereas *β* controls the curvature at lower means. We fitted these two parameters by hand. Their interpretation is that, 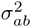 corresponds to the minimal amount of cell-to-cell variation in the product *ab* that is possible for any promoter architecture. The variable *β* = 450 roughly corresponds to the burst-size.

**Figure S15.**
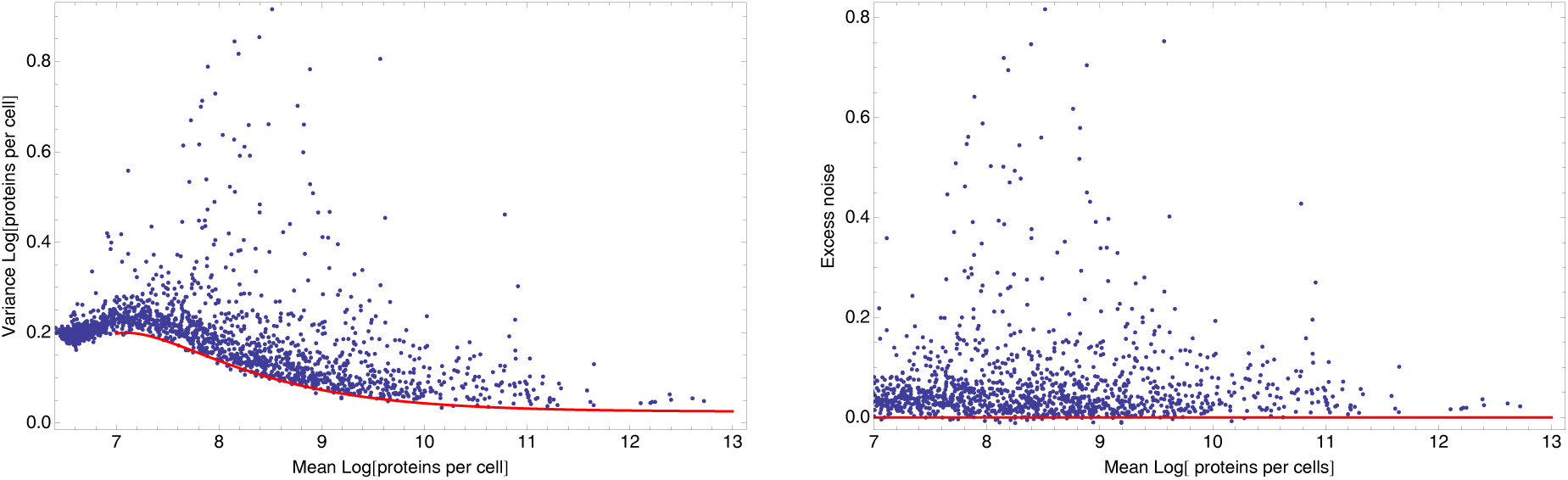
Dependence between mean and variance of log FACS intensities. **Left panel:** Means and variances of log-FACS intensities of all native promoters (blue dots) together with a fitted lower bound on the variance as given by equation (66) using 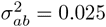 and *β* = 450 (red curve). **Right panel:** Excess noise (obtained by subtracting the fitted lower bound from the variance) as a function of mean log-FACS intensity for all native promoters (blue dots). The red line shows the *x*-axis.

Note that the log-fluorescence on the horizontal axis corresponds to the sum of fluorescence resulting from GFP molecules and the background fluorescence. The estimated background level *n*_bg_ = 582.3 corresponds to a log-fluorescence of 6.37. The region on the horizontal axis between 6.37 and 7 ≈ log(2 * 582.3), thus corresponds to cells where the fluorescence due to GFP molecules is less than the background fluorescence. In this regime the noise distribution results from a combination of fluctuations in background fluorescence and in protein numbers (which may be correlated because part of these fluctuations may result from fluctuations in cell size) and our noise model (66) breaks down. In the following we will focus on those promoters with fluorescence due to GFP at least as large as the background fluorescence, i.e. with mean log-fluorescence larger than 7.

To obtain a deviation of each promoter's variance from the minimal variance that is possible at its expression level, we define the *excess noise η* as the difference between a promoter's variance and its the fitted minimal variance 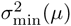 as given by equation (66) with *β* = 450 and 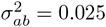:

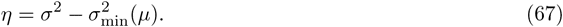

The right panel of **figure S15** shows the excess noise levels of all native promoters as a function of their means. The figure shows that, with this correction, there is no longer any systematic dependence between mean expression levels and noise. Therefore, we can use excess noise as a measure of transcriptional noise that allows us to compare noise levels of promoters with different mean expression levels.

### FACS Selection

As explained in the **Methods**, for both the medium and high expression evolutionary runs, the desired expression level *µ_∗_* is taken from the expression level of a reference promoter from the library of E. coli promoters. At each selection round we measure the expression *µ_∗_* of the reference promoter, and set the center of the FACS's selection window to *µ_∗_*. We then set the width of the selection window such that 5% of the cells have expression levels within the selection window.

Although, in principle, the FACS's selection should work such that a cell with expression level anywhere within the selection window has 100% probability to be selected, and 0% probability to be selected if the cell's expression is anywhere outside the selection window, it is unrealistic to assume that the boundaries of the selection window are so precisely defined in practice. As illustrated below, comparison of the population's expression levels before and after selection shows that the probability for a cell with log-fluorescence *x* to be selected can be well-approximated as

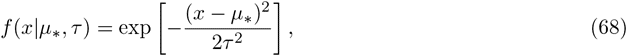

where *µ_∗_* is the desired expression level and *τ* corresponds to the width of the selection window.

Note that, for a promoter with mean expression *µ* and variance *σ*^2^, the fraction *P* (*x µ, σ*) of its cells that have expression level *x* is given by

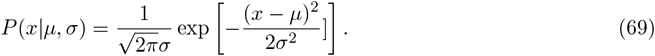

Consequently, the ‘fitness’ of this promoter, i.e. the fraction of its cells that are selected in the FACS, is given by

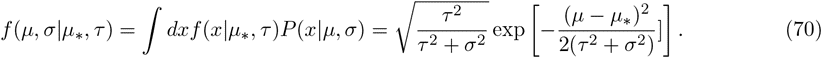

To infer the values of *µ_∗_* and *τ* that apply to our evolutionary runs, we performed a number of experiments in which we:

1. Took a population from one of the rounds of our evolutionary runs.
2. Measured its distribution of log-fluorescence levels.
3. Set the selection window [*µ_∗_ δ, µ_∗_* +*δ*] such that a percentage *p* of the population has log-fluorescence levels within this selection window.
4. Performed selection and re-measured the log-fluorescence levels of the selected population.

As shown in **Fig. 1b** of the main paper, in the evolutionary runs which are selecting for high expression, the selection window is changing at every round of the evolutionary run. In contrast, in the evolutionary runs selecting for medium expression, the selection window is essentially constant from round 3 through round 5 of the run. We thus decided to focus on inferring the precise fitness function that acted during these 3 rounds of selection.

We took the evolved populations from the third and fifth round of the evolutionary runs selecting for medium expression, and performed another round of selection on them, selecting 5% of the population closest to the desired log-fluorescence *µ_∗_*. In addition, we also performed a round of less stringent selection on these populations, selecting 25% closest to the desired level, and a round of more stringent selection, selecting only the 1% of the population closest to the desired level. Besides measuring the log-fluorescence levels of the population both before and after the round of selection, we also selected dozens of clones from the populations before and after the selection, and measured the entire distribution of log-fluorescence **Figure S16** shows the means and variances of the log-fluorescence distributions of these clones.

**Figure S16.**
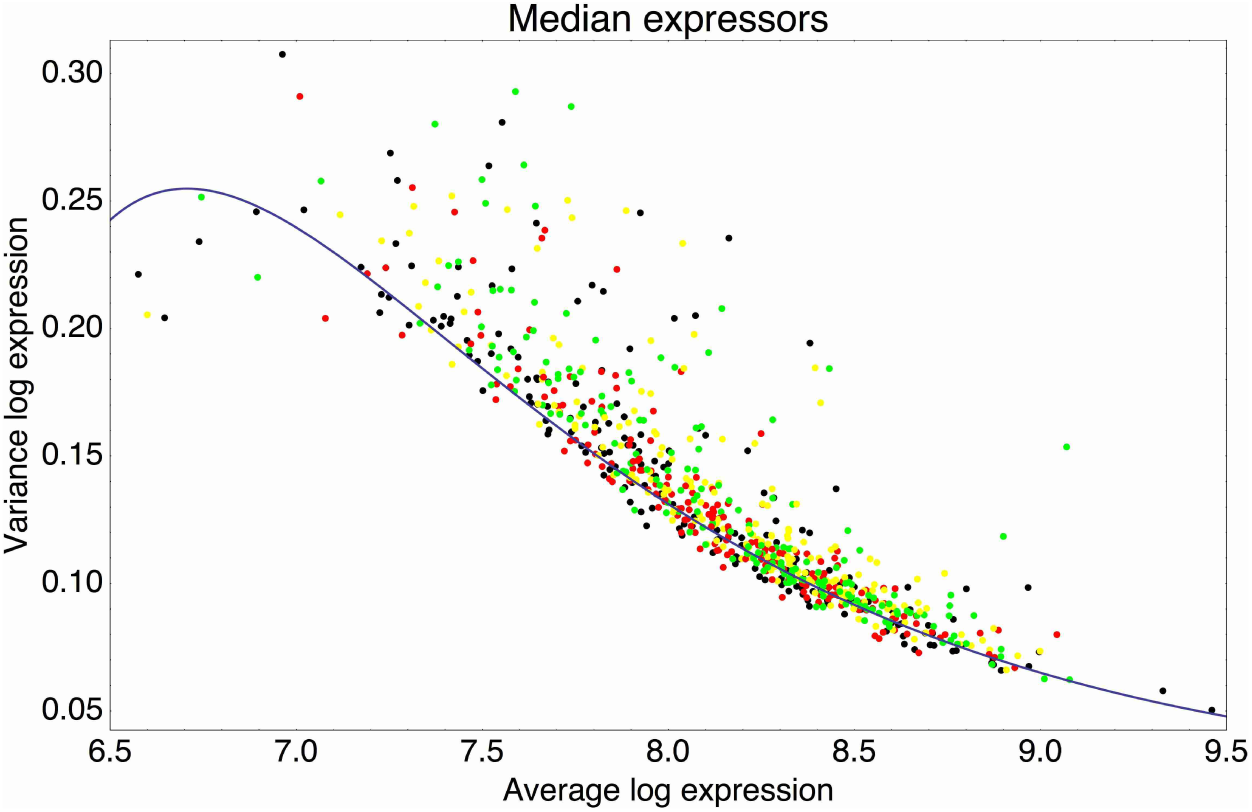
Means and variances of the log-fluorescence levels of clones from the third and fifth rounds of the evolutionary runs in which we selected for medium expression (black dots), and clones obtained after performing another round of selection on these populations, selecting either 1% (red), 5% (yellow), or 25% of the population closest to the desired log-fluorescence µ∗. The blue curve shows an approximate fit of the typical variance σ^2^ as a function of the mean *µ*: *σ*^2^(*µ*) = 0.02 + 384*e*^−*µ*^ − 156′915*e*^−2*µ*^. levels for these clones.

Intuitively, one might think that the relative fitness *f* (*x*) of each log-fluorescence level *x* could be easily estimated by simply measuring the ratio of the fraction of the population *p*′(*x*) with log-fluorescence level *x* after selection and the fraction *p*(*x*) with log-fluorescence *x* before selection. However, the single cells that were selected in the FACS each grow into an entire population of cells before the ‘after selection’ population is measured again. Thus, a selected cell containing a promoter with a given mean *µ* and variance *σ*^2^ will contribute an entire population of cells with this distribution, even though the individual cell may itself have had a log-fluorescence that was in one of the tails of this distribution. Thus, in general the distribution of log-fluorescence levels in the population after selection may be much wider than the actual selection window itself.

Before selection, the population consisted of a mixture containing (unknown) fractions *ρ*(*µ, σ*) of cells containing promoters with mean *µ* and variance *σ*^2^. This gives rise to an overall distribution *p*(*x*) of expression levels given by

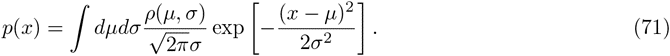

Unfortunately we cannot uniquely infer *ρ*(*µ, σ*) from knowing only the distribution *p*(*x*). However, as shown in **Fig. S16**, for the clones in these populations, the large majority of promoters have variances *σ*^2^ lying in a narrow band as a function of *µ*. We thus chose to make the approximation that *all* promoters in the population have variances *σ*^2^ that are uniquely determined by their mean expression *µ*, and we used the fit *σ*^2^(*µ*) shown as the blue curve in **Fig. S16**. This simplifies the problem from inferring a two-dimensional distribution *ρ*(*µ, σ*) to inferring a one-dimensional distribution *ρ*(*µ*), i.e.

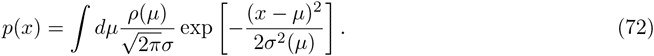

Applying selection with target log-fluorescence *µ_∗_* and width *τ* to this distribution, we obtain a new distribution

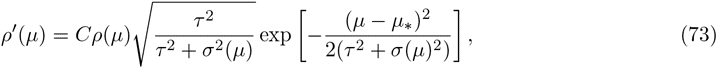

where *C* is a normalization constant that ensures 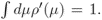. The population distribution of log-fluorescence levels after selection is then

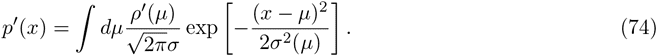

To infer the parameters of the fitness-function for each selection that we performed, we fit the distribution *ρ*(*µ*), and parameters *µ_∗_* and *τ* that lead to an optimal fit to the observed distributions *p*(*x*) and *p*′(*x*). **Figure S17** shows the inferred and observed distributions, as well as the inferred fitness function, for each of the 6 selection experiments that we performed.

**Figure S17.**
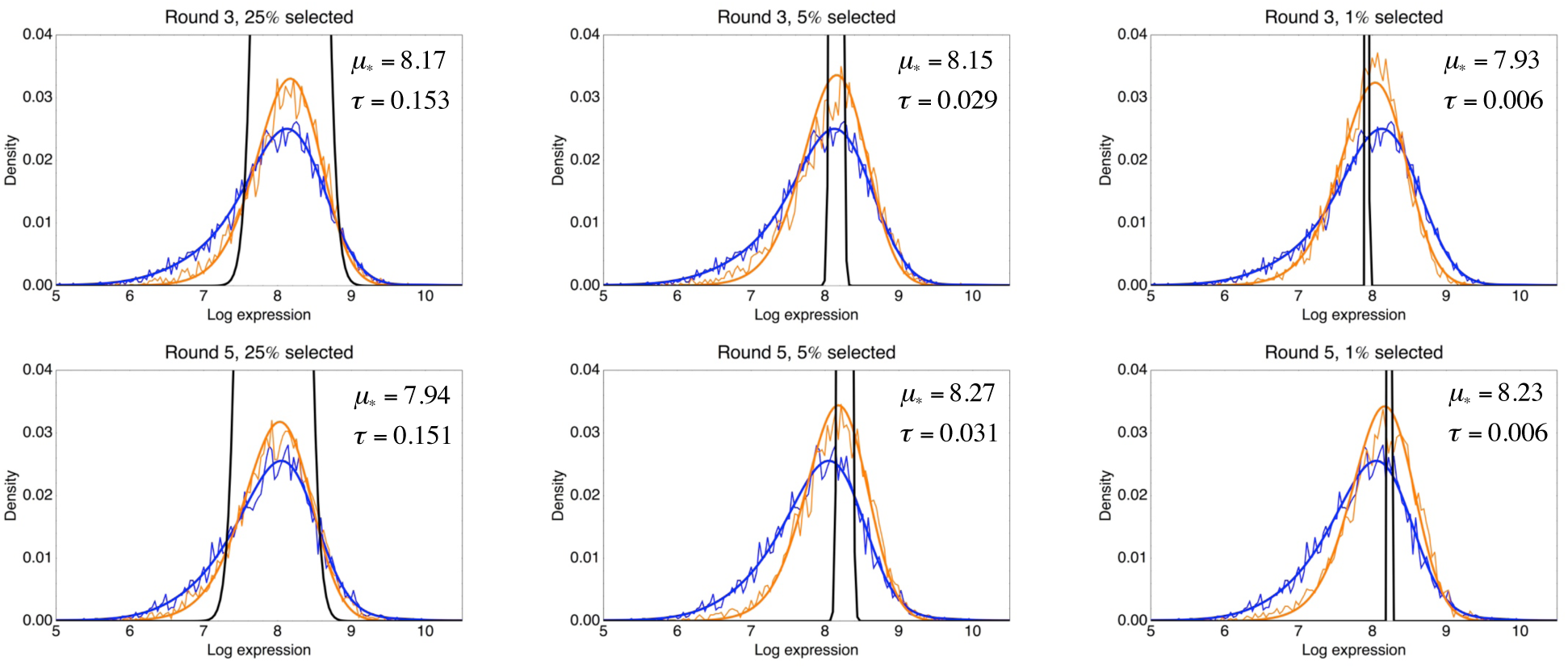
Inference of the fitness function from the observed log-fluorescence distribution before and after a round of selection. Each panel corresponds to one selection experiment with the title indicating on which population an extra round of selection was performed, i.e. a population either from the third or fifth round of the evolutionary run for medium expression. The thin blue line indicates the observed log-fluorescence distribution *p*(*x*) before selection, and the thin orange line the observed distribution *p*′(*x*) after selection. The thick lines show the corresponding fitted distributions. The inferred selection window *f*(*x*|*µ*∗, *τ*), i.e. equation (68), is indicated in black, and its parameters *µ*_∗_ and *τ* are indicated in each panel as well.

The figure shows that the distributions *p*(*x*) and *p*′(*x*) can be well fit by this model, illustrating that the form of the selection window, equation (68), can well describe the effects of selection in the FACS machine. Moreover, we see that the distributions *p*(*x*) and *p*′(*x*) are typically significantly wider than the selection window *f*(*x|µ_∗_, τ*). Moreover, the fitted values of *τ* are almost perfectly proportional to the fraction of the population that was selected, with a value of *τ* ≈ 0.03 corresponding to the selection of 5% of the population that was used during the evolutionary runs. The fits also show that, although we in each experiment determine *µ_∗_* from the expression of the same reference promoter, there is some variability in *µ_∗_* from one experiment to the next. From these 6 experiments, we find that on average 〈*µ_∗_*〉 = 8.115 and *σ*(*µ_∗_*) = 0.133.

Thus, in each selection round the fitness of a promoter with mean *µ* and variance *σ*^2^ is given by expression (70), where *τ* = 0.03 and *µ_∗_* fluctuates around 〈*µ_∗_*〉 = 8.115. The effective fitness experienced by this promoter is thus given by the geometric average of equation (70) with fluctuating values of *µ*_∗_ and this is given by

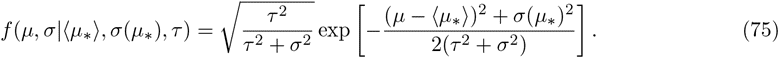

**Figure S18** shows a contour plot of this fitness function with the inferred parameters of 〈*µ_∗_*〉, *σ*(*µ_∗_*) and *τ* as a function of the mean fitness *µ*, and the excess noise level *η* = *σ*^2^ − *σ*^2^_min_ (*µ*), where 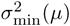 is the minimal variance as a function of mean expression level *µ*, equation (66). Note that for these measurements the plasmids were transformed into a different strain than those used to compare with the native E. coli promoters. We noticed that the minimal noise level 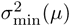 as a function of mean *µ* is slightly different. Although the background fluorescence and burst-size parameter *β* = 450 are the same, the parameter 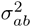 is smaller, i.e. 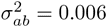 instead of 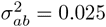.

**Figure S18.**
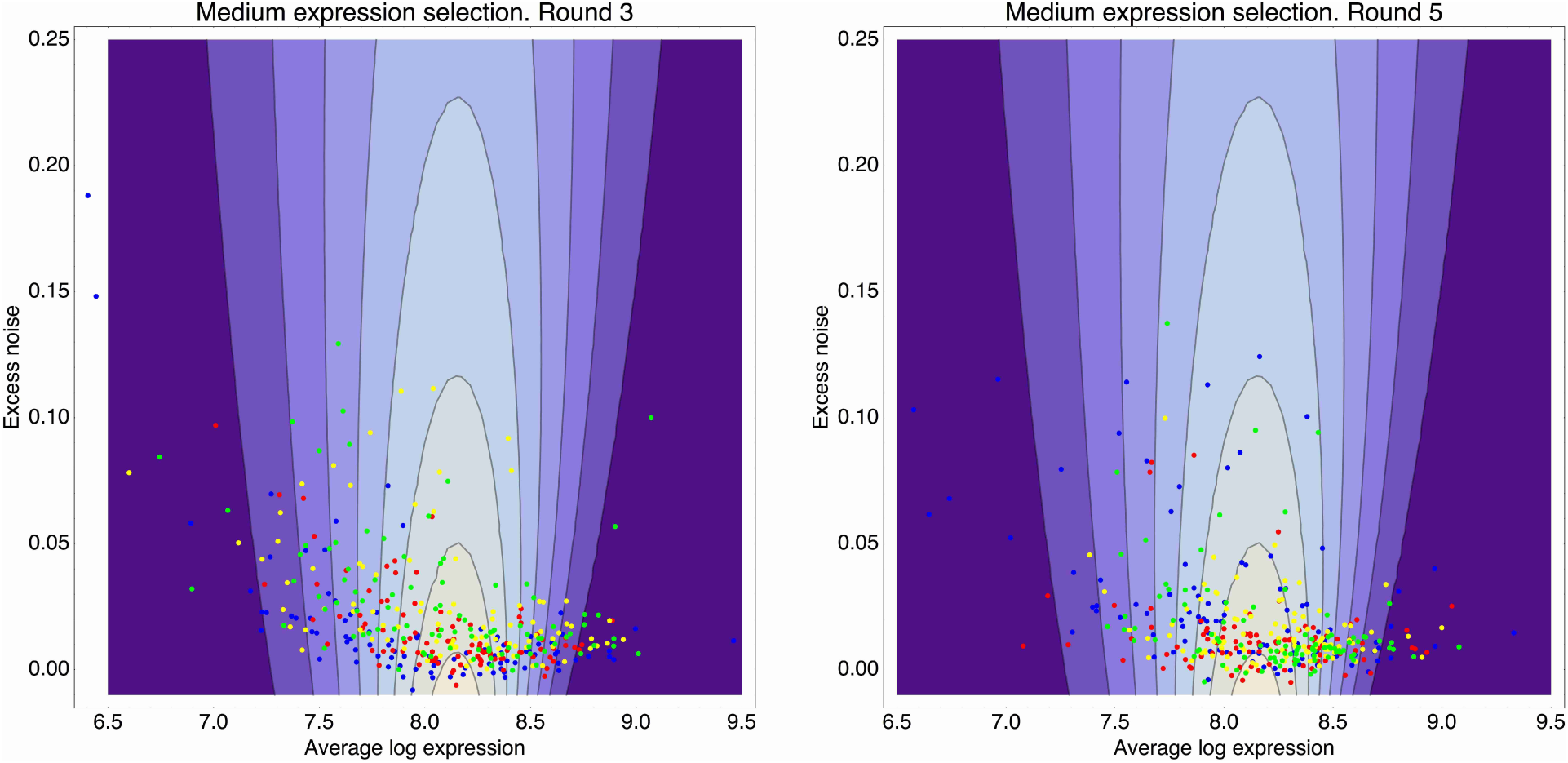
Contour plot of the inferred fitness function (75) as a function of mean expression *µ* (horizontal axis) and excess noise (vertical axis), that acts on the population from rounds 3 through 5 of the evolutionary runs for medium expression. The contours correspond to fitness values (fraction of cells selected) of 0.01, 0.02, 0.03, through 0.08. **Left panel:** In addition to the fitness function (contours) the panel shows the means and excess noise levels of a selection of clones from the third round of the evolutionary run (blue dots), and clones that resulted from subjecting this population to another round of selection, selecting either for the 1% (red dots), 5% (yellow dots), or 25% (green dots) of cells with expression closest to the desired expression level. **Right panel:** As in the left panel, but with the dots corresponding to clones from the 5th round of the evolutionary run, and clones resulting from additional rounds of selection on this population (colors as in the left panel).

**Figure S18** clearly shows that fitness drops far more dramatically as a function of mean *µ* than as a function of excess variance *η*, i.e. except for right at the optimal mean *µ_∗_*, the contours are running almost vertically in the plot. We do not observe promoters with high excess noise, even though their fitness would easily allow it. For example, a promoter with mean expression near the optimum 8.11 but excess noise as high as 0.25 (i.e. significantly higher than observed for any of the clones) would have higher fitness than any promoter with mean less than *µ* = 7.76 or larger than *µ* = 8.47 (independent of their noise), even though we observe many promoters with means that deviate this far from the optimum.

The fact that we do not observe high excess noise promoters, even though they would not be selected against, strongly suggests that such high noise promoters are uncommon *a priori*, i.e. among all random sequences that drive expression at a medium level, the large majority have low excess noise levels and high noise promoters are rare. Moreover, note also that for promoters that are not near the optimum, the optimal excess noise level is typically *larger* than those of the observed clones, e.g. the optimal excess noise for a promoter with mean *µ* = 7.5 is *η* = 0.22. These observations all suggest that promoters have not experienced significant selection on their noise levels.

To further support this conclusion, **Fig. S19** shows the inferred fitness values for the observed clones both as a function of their mean expression (left panel) and as a function of their excess noise (right) panel. The figure shows that, whereas the fitness of a clone can be accurately predicted from its mean *µ*, fitness is almost entirely uncorrelated to a promoter's excess noise *η*.

Finally, if there was significant selection on noise levels, then we expect noise levels to systematically shift under selection. **Figure S20** shows cumulative distributions of excess noise levels for clones obtained from different populations of cells.

**Figure S19.**
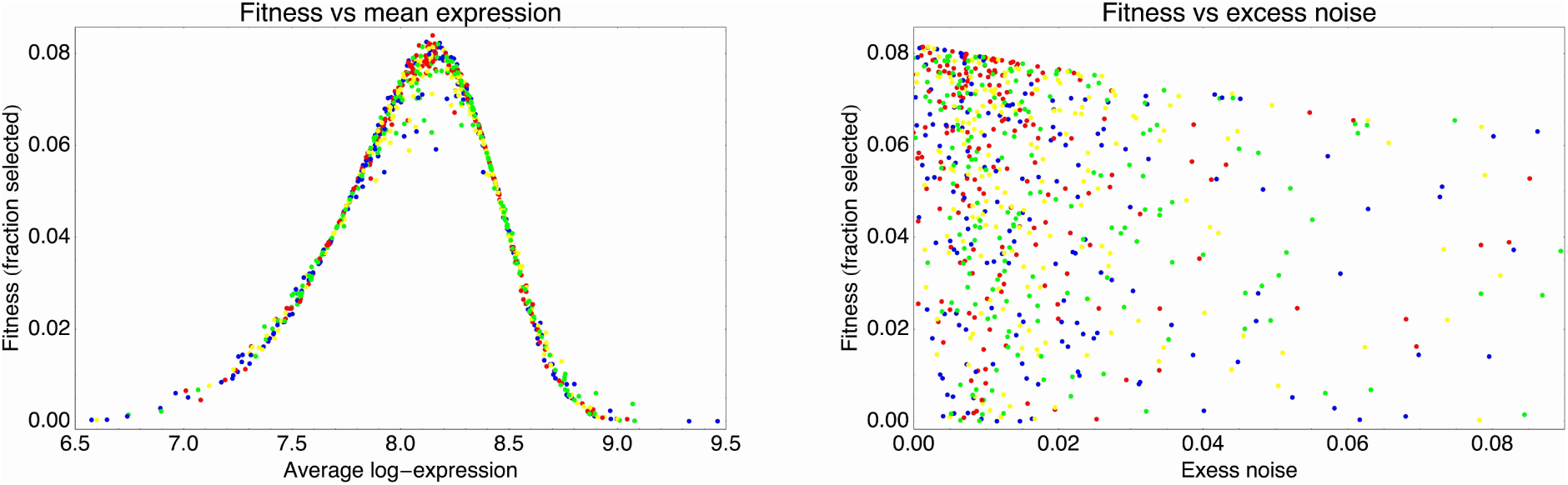
Fitness of the observed clones, as given by equation (75), as a function of their mean expression *µ* (left panel) and their excess noise *η* (right panel). As in figure S18, the blue dots correspond to clones from the third and fifth round of the evolutionary run, the red dots result from another round of stringent selection (top 1%), the yellow dots from another round of standard selection (top 5%), and the green dots from a round of weaker selection (top 25%).

**Figure S20.**
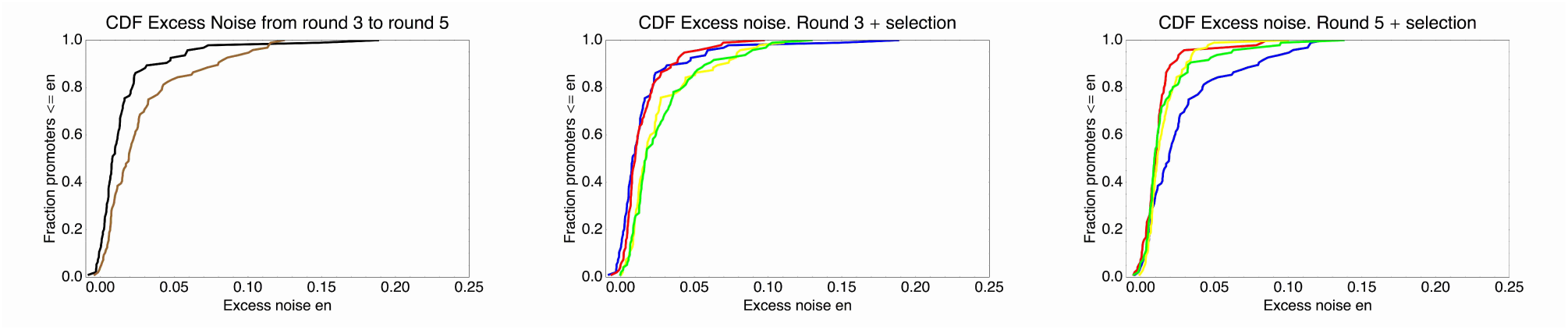
Cumulative distribution functions of excess noise levels for the promoters extracted from different populations. **Left panel**: Excess noise levels of promoters from the 3rd (black) and 5th (brown) round of the evolutionary run. **Middle panel**: Excess noise levels of promoters from the 3rd round of the evolutionary run (blue), and from clones that resulted from another round of either stringent (red), normal (yellow), or weak (green) selection. **Right panel**: As in the middle panel but now for clones from the 5th round and clones resulting from another round of selection on this population.

The figure shows that, surprisingly, the excess noise levels seem to increase from the third to the fifth round in the evolutionary run. However, given the limited number of clones involved, the change in excess noise levels is only marginally significant (*p* = 0.004 in a t-test). Similarly, the effect of selection on excess noise levels seems to be opposite on the populations of round 3 and round 5 (center and right panels in **Fig. S20**). We suspect that there are some systematic experimental fluctuations that make measured excess noise levels vary across days, and that the observed distributions of excess noise levels are more a reflection of experimental variability than of true shifts in the distribution. Importantly, excess noise levels larger than 0.1, which are observed for a substantial fraction of native promoters, are very rare for all these populations.

In summary, our in depth analysis of the FACS fitness function and the effects of selection show that the noise properties of the synthetic promoters have not been significantly shaped by selection. Already the synthetic promoters at the third round of selection are tightly concentrated in a low noise band, even though selection does not select against low noise, and promoters with mean away from the optimum would benefit from having higher noise. Two additional rounds of mutation and selection on the promoters from the third round do not substantially change the distribution of noise levels confirming that there are no substantial fitness differences among promoters with different noise. Similarly, performing additional rounds of selection (be it very stringent, normal, or lenient) also do not substantially change the observed noise levels of the selected promoters. Thus, our results show that promoters selected from a large collection of random sequences naturally display low noise levels. Importantly, this implies that the native promoters with substantially higher noise levels must have experienced some selective pressures that caused them to increase their noise.

### A simple model for the evolution of gene regulation and expression noise

Given a particular environment, the fitness, e.g. growth-rate or survival probability of a cell, depends on the expression level of its genes. Note that the fact that gene regulatory mechanisms have evolved already demonstrates that different environments require different gene expression patterns, i.e. expressing a gene at the ‘wrong’ level for a given environment has negative effects on fitness/growth-rate. For simplicity, we will focus on a single gene. We assume that, in a given environment, there is an optimal expression *µ_∗_* level. Given that, as we have seen, expression levels are roughly log-normally distributed, we will express expression levels in log-space, i.e. the logarithm of the number of proteins per cell. We define that fitness at the optimal expression level *µ_∗_* as *f_o_*. Fitness will fall as the expression level *x* moves away from this optimum. In this simple conceptual model, we will assume that, like in our FACS selection, the fitness *f*(*x*) falls approximately Gaussian away from the optimum, i.e.

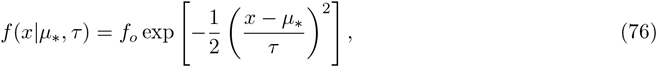

where *τ* is again a parameter that determines how fast the fitness falls when the expression *x* moves away from the optimum *µ_∗_*.

To justify the Gaussian form of the fitness function, assume that the fitness is determined by the growth of the population over some characteristic time *t*. That is, if cells grow at rate *ρ*, then the fitness is *f* = *e^ρt^*. The growth rate *ρ* is optimal at *x* = *µ_∗_*, and to second order in the difference between *x* and *µ_∗_*, we can write

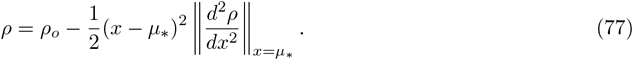

Defining *f_o_* = *_e_^ρt^* and 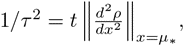 we obtain the fitness function defined above.

In complete analogy with the FACS selection case, the fitness *f*(*µ, σ|µ_∗_, τ*) of a promoter with mean *µ* and variance *σ*^2^ in an environment with optimal expression level *µ_∗_* and width *τ* is given by the integral of the product of the fitness function (76) and the Gaussian distribution of expression levels, giving

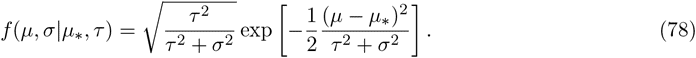

Note that this functional form is a reasonable approximation to the fitness function as long as expression levels are roughly log-normally distributed, and as long as the integral of expression levels and fitness function can be approximated using the standard Laplace approximation, i.e. expanding the logarithm of fitness to second order around its maximum.

We now extent this simple situation in two respects. First, instead of assuming that the optimal level *µ_∗_* is fixed, we imagine that the population of cells has gone through several different ‘environments’, where in each environment *e* there was an optimal expression level *µ_e_*. For simplicity we assume *τ* is the same in each environment.

Let's first consider what this situation implies for the fitness of a promoter expressing at mean level *µ* with variance *σ*^2^. The number of offspring that a strain with mean *µ* and variance *σ*^2^ produces (or leaves behind) after experiencing environment *e* is proportional to *f*(*µ, σ|µ_e_, τ*). Consequently, the final number of offspring produced after experiencing all environments is given by the product 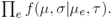 We define the overall log-fitness log[*f*(*µ*, *σ*)] as the average of the log-fitness across all environments:

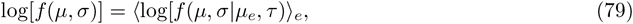

where the subscript *e* indicates that we are averaging over all environments *e* (which we drop for convenience from here on). Using the expression (78) we obtain

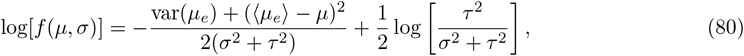

where 〈*µ_e_*〉 is the average ‘desired’ expression level, and var(*µ_e_*) is the variance in desired expression levels across the environments. It is immediately clear from equation (80) that, as a function of the mean expression *µ*, optimal fitness is obtained when *µ* = 〈*µ_e_*〉. Substituting this optimal mean level, we find that optimal variance is given by

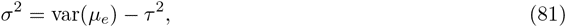

when var(*µ_e_*) ≥ *τ*^2^, and *σ* = 0 otherwise. That is, when the variance in desired expression levels is larger than the width of the selection window *τ*, then a strain can increase its fitness by raising the noise-level *σ* of the promoter. This result is equivalent to results on selection for phenotypic variance obtained previously, e.g. [10, 23]. However, in these previous models that more abstractly considered ‘phenotypic traits’, it was assumed that both the mean and variance of the phenotypic trait were not only directly encoded by the genotype, but could also be independently altered through mutations, without explicitly considering how mean and variance would be encoded in the genotype. In our case, where the ‘trait’ under study is the transcription rate of a promoter, it is a priori quite clear how mutations may alter mean levels, e.g. through changes in the affinity of the sigma-factor binding site, but much less clear how the variance is encoded in the genotype. Moreover, rather than simply increasing its noise, we would naturally expect that promoters would evolve *gene regulation* in order to deal with different required expression levels across different environments.

### Including gene regulation

We now further extend the model by considering that there are various transcriptional regulators in the cell whose activities may vary across the different environments *e*. By evolving binding sites for a transcription factor, the promoter becomes regulated by it and, consequently, the mean expression *µ* becomes a function *µ*(*e*) of the environment *e*.

For simplicity we consider the case of a single regulator whose mean activity (i.e. concentration of the DNA-binding version of the regulator) *r_e_* is a function of the environment *e*. Since the transcriptional regulator's expression will itself also be subject to gene expression noise, the activity of the regulator varies from cell to cell. We will assume that, in each environment, the activity of the regulator varies from cell-to-cell in a roughly Gaussian manner, with variance 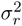, i.e. the probability to find a cell in environment *e* with regulator level *r* is

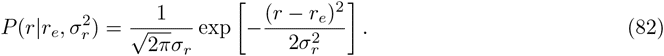

We characterize the regulation of the promoter by the regulator through a single *coupling constant c* such that, in cells with regulator level *r*, the distribution of expression levels is a Gaussian with mean *µ* + *cr* and variance *σ*^2^, i.e.

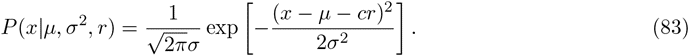

Integrating over the distribution of regulator levels *P* (*r|r_e_, σ*^2^), the final distribution of expression levels is given by a Gaussian with mean *µ*(*e*) = *µ* + *cr_e_* and variance *σ*^2^ + *c*^2^*σ*^2^, i.e.

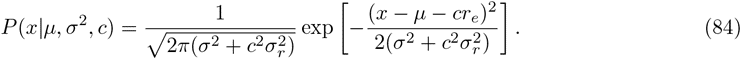

In environment *e*, with desired level *µ_e_*, the fitness of a promoter with coupling *c* is then given by

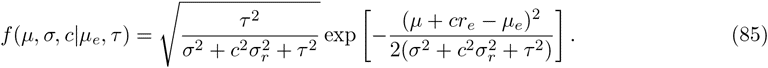

The log-fitness, averaged over all environments *e*, is given by

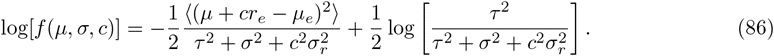

It is again easy to see that, with respect to the mean expression *µ*, fitness is optimized when *µ* = 〈*µ_e_*〉 − *c*〈*r_e_*〉. In the following we will assume that the mean expression *µ* matches this optimal value.

We can rewrite the expression for the average log-fitness in a simpler form by introducing the following set of effective parameters. First, the variable

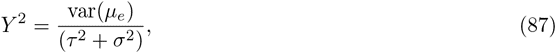

measures the variance in desired expression levels *µ_e_* relative to the sum of the variances associated with the width of selection *τ* ^2^ and the noise of the unregulated promoter *σ*^2^. The variable *Y* quantifies the ‘expression mismatch’ between the promoters average expression *µ* and the (varying) desired expression levels *µ_e_*. The second effective parameter

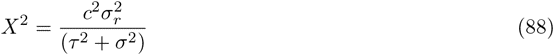

measures the strength of the regulator coupling constant *c*. More precisely, it quantifies the contribution *c*^2^*σ*^2^ to the promoter's variance in gene expression, again relative to (*σ*^2^ + *τ* ^2^). We will refer to *X* as the coupling constant. Third, the parameter

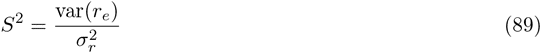

measures the ‘signal-to-noise’ ratio of the regulator, i.e. the variance var(*r_e_*) of its mean level across conditions, relative to its variance *σ*^2^ within each condition. A regulator with large *S* varies a lot in activity across environments and has relatively little noise in each, whereas a regulator with small *S* varies little across environments relative to its noise level. Finally, we have the correlation *R* between the desired expression levels *µ_e_* and the regulator's activities *r_e_*, i.e.

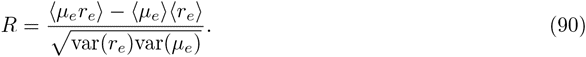

In terms of these parameters we have for the average log-fitness

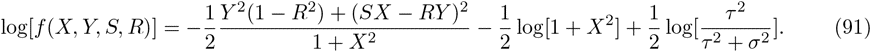

The last term is a constant that does not depend on our effective parameters and we will ignore it from now on.

We intuitively expect that the promoter's fitness would benefit most from coupling to a regulator that is perfectly correlated with the environment's requirements, i.e. at *R* = 1. Indeed we find that the derivative log[*f*(*X*)] with respect to *R* is given by

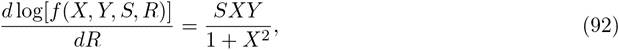

which is positive as long as the desired levels vary (*Y* > 0), the regulator has some variation across environments (*S* > 0) and there is positive coupling (*X* > 0). Thus, in general, if we keep all other variables fixed, an increase in the regulator's correlation *R* is always beneficial.

We now consider the case in which a regulator with a given correlation *R* and signal-to-noise rate *S* is given, and we want to determine the optimal coupling *X_∗_* that maximizes log[*f*(*X, Y, S, R*)] as a function of the expression mismatch *Y*. The derivative of log[*f*(*X, Y, S, R*)] with respect to *X* is given by

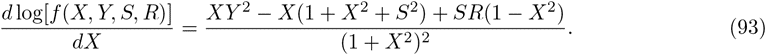

At *X* = 0, this derivative equals *SR*. Thus, whenever *R* > 0, the derivative is positive at *X* = 0. Because, as can be easily seen from equation (93), the derivative is guaranteed to be negative for large *X*, this implies that, whenever *R* > 0, there is an optimal coupling *X_∗_* that is positive, i.e. *X_∗_* > 0. Thus, whenever *R* > 0, the promoter is guaranteed to increase its fitness by evolving a nonzero coupling to the regulator.

The optimal coupling *X_∗_* is given by the positive solution of the third order polynomial in the numerator of (93). In general we find that, when *Y* is small, the optimal coupling is given by

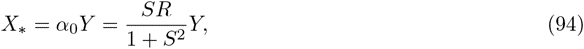

and that when *Y* is very large *X_∗_* obeys

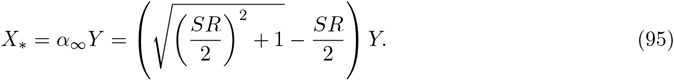

That is, both for very small and very large *Y*, the optimal coupling *X_∗_* is directly proportional to *Y*, with proportionality constants *α*_0_ and *α_∞_*, respectively. Moreover, *α_∞_* ≥ *α*_0_. The behavior of *X_∗_* as a function of *Y*, for different values of *R* and *S* is illustrated in **Fig. S21**.

**Figure S21.**
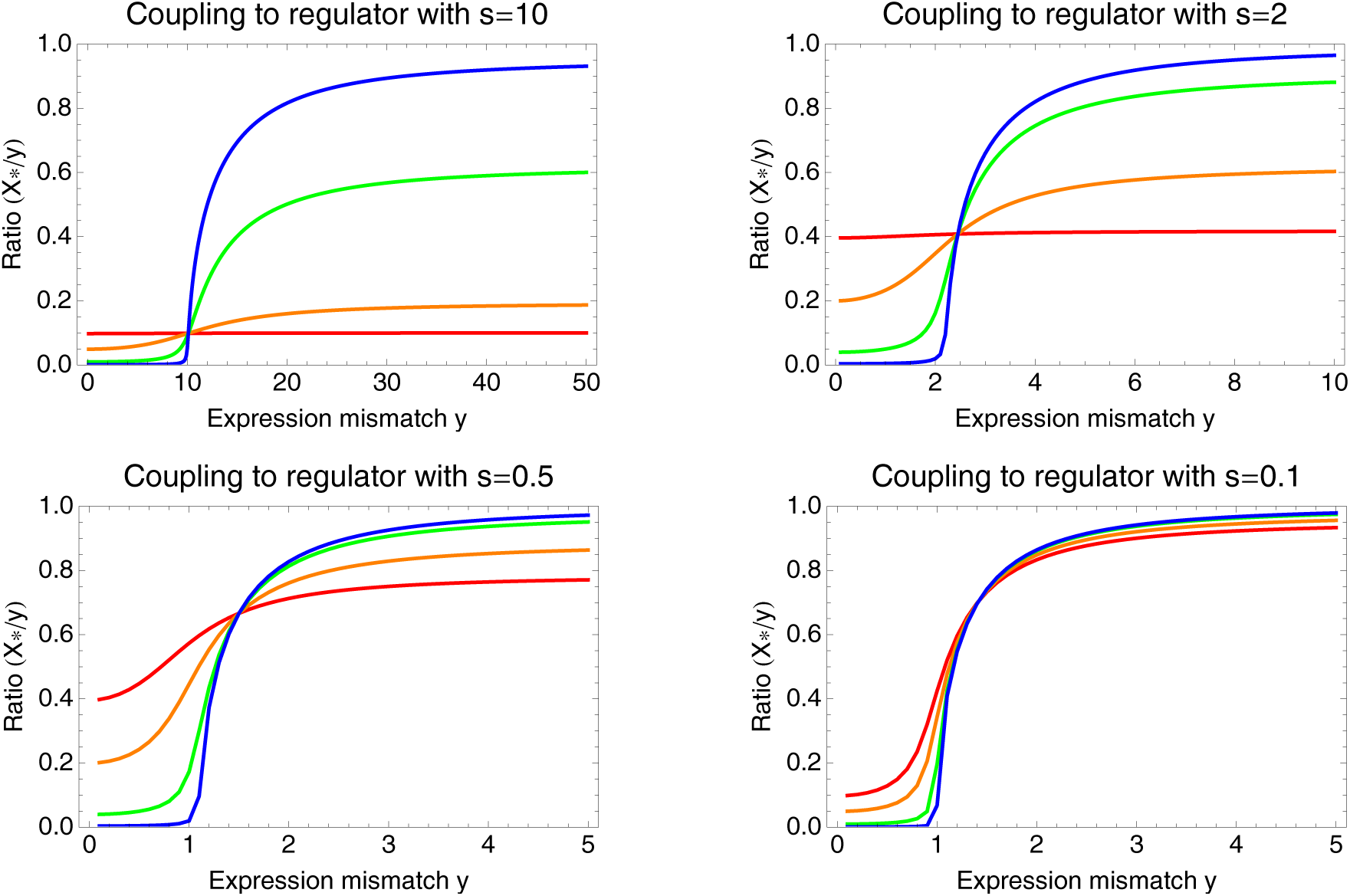
The ratio *X*_∗_/*Y* between optimal coupling *X*_∗_ and expression mismatch *Y* as a function of *Y*,for different values of the regulator's signal to noise ratio *S* and the correlation between regulator and environment *R*. Each panel corresponds to a different signal to noise ratio *S*, from a high signal regulator in the top left, to a noisy regulator at the bottom right. In each panel, the different colored lines correspond to different correlations *R*, i.e. *R* = 0.01 (blue), *R* = 0.1 (green), *R* = 0.5 (orange), and *R* = 0.99 (red).

The figure shows that, as *Y* increases the ratio *X_∗_/Y* switches from the lower *α*_0_ to the higher *α_∞_*. Whenever both the correlation *R* and the signal-to-noise *S* are high (the orange and red curves in the top two panels), there is only a small difference between *α*_0_ and *α_∞_*. That is, *X_∗_* increases roughly linearly with *Y* when there is a well-correlated regulator with high signal-to-noise.

In contrast, when the correlation *R* is low or the regulator is noisy, there is a large difference between *α_∞_* and *α*_0_. Moreover, the optimal coupling shows a sharp transition from low values to much higher values at a ‘critical’ value of *Y*. This critical value of *Y* occurs at 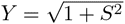 when *R* is low (the blue and green curves), and slightly earlier when *R* increases (orange and red curves). When the regulator is very noisy (bottom right panel of **Fig. S21**) the behavior of *X_∗_* becomes almost independent of the correlation *R*, showing a sharp transition from almost no coupling to strong coupling when *Y* ≈ 1. This behavior even extends to the case where there is no correlation whatsoever between the regulator and the environment, i.e. *R* = 0.

### Coupling to an uncorrelated regulator

When *R* = 0 the optimal coupling is given by

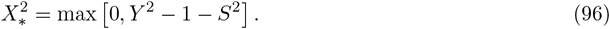

That is, at the critical value 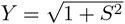 the coupling goes from zero to a positive value. For large *Y* the optimal coupling is simply *Y*. The behavior of optimal coupling *X_∗_* as a function of *Y* is shown in **figure S22**.

**Figure S22.**
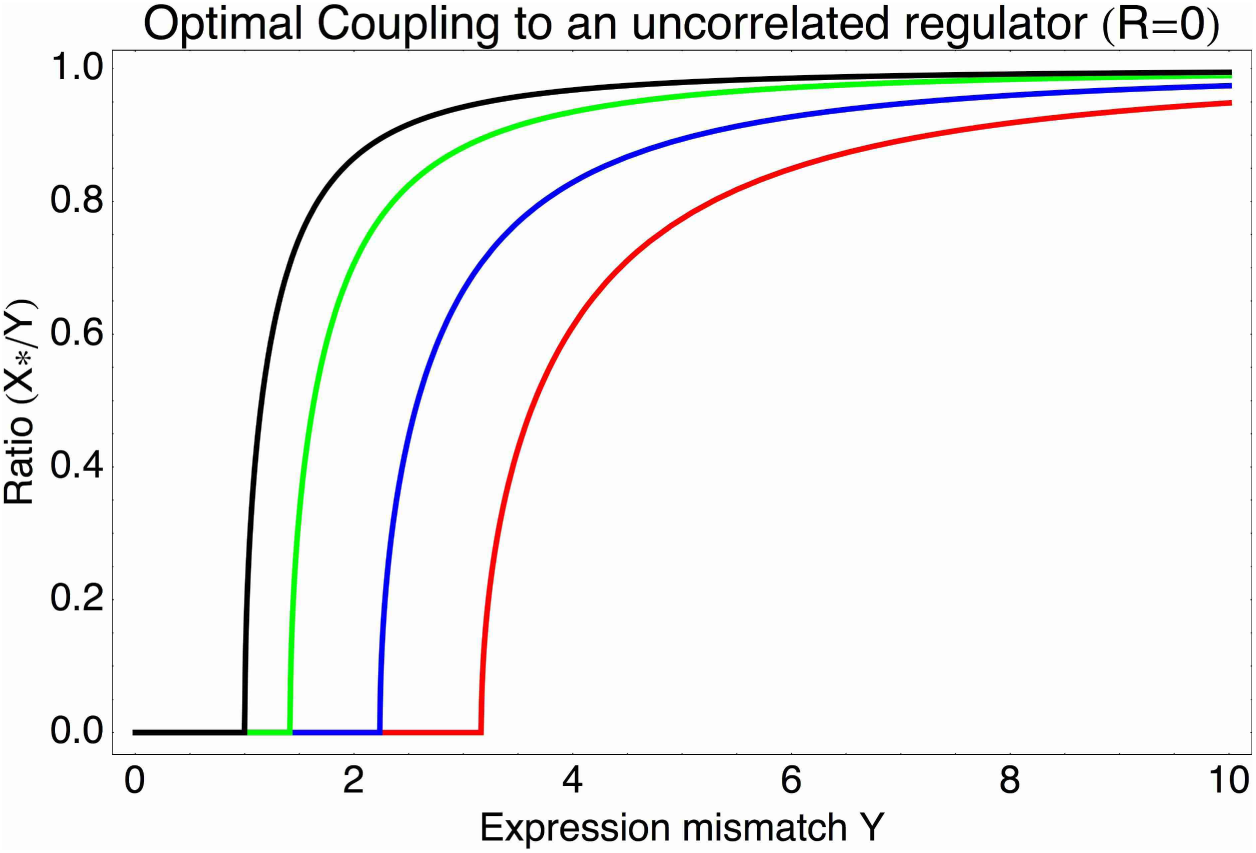
Optimal coupling *X*_∗_ as a function of the expression mismatch *Y* for different values of the signal-to-noise ratio *S*, i.e. *S* = 0 (black), *S* = 1 (green), *S* = 2 (blue), and *S* = 3 (red).

### Log-fitness at optimal coupling *X*_∗_

We next consider the case in which the promoter has a certain expression mismatch *Y*, and we calculate the log-fitness that it can obtain by *optimally* coupling to a regulator that has a certain signal-to-noise *S* and correlation *R*. **Figure S23** shows the resulting log-fitness values as a function of *S* and *R* for 4 different values of the expression mismatch *Y*: *Y* = 1 in the top-left panel, *Y* = 2 in the top-right panel, *Y* = 4 in the bottom left panel, and *Y* = 8 in the bottom right panel.

**Figure S23.**
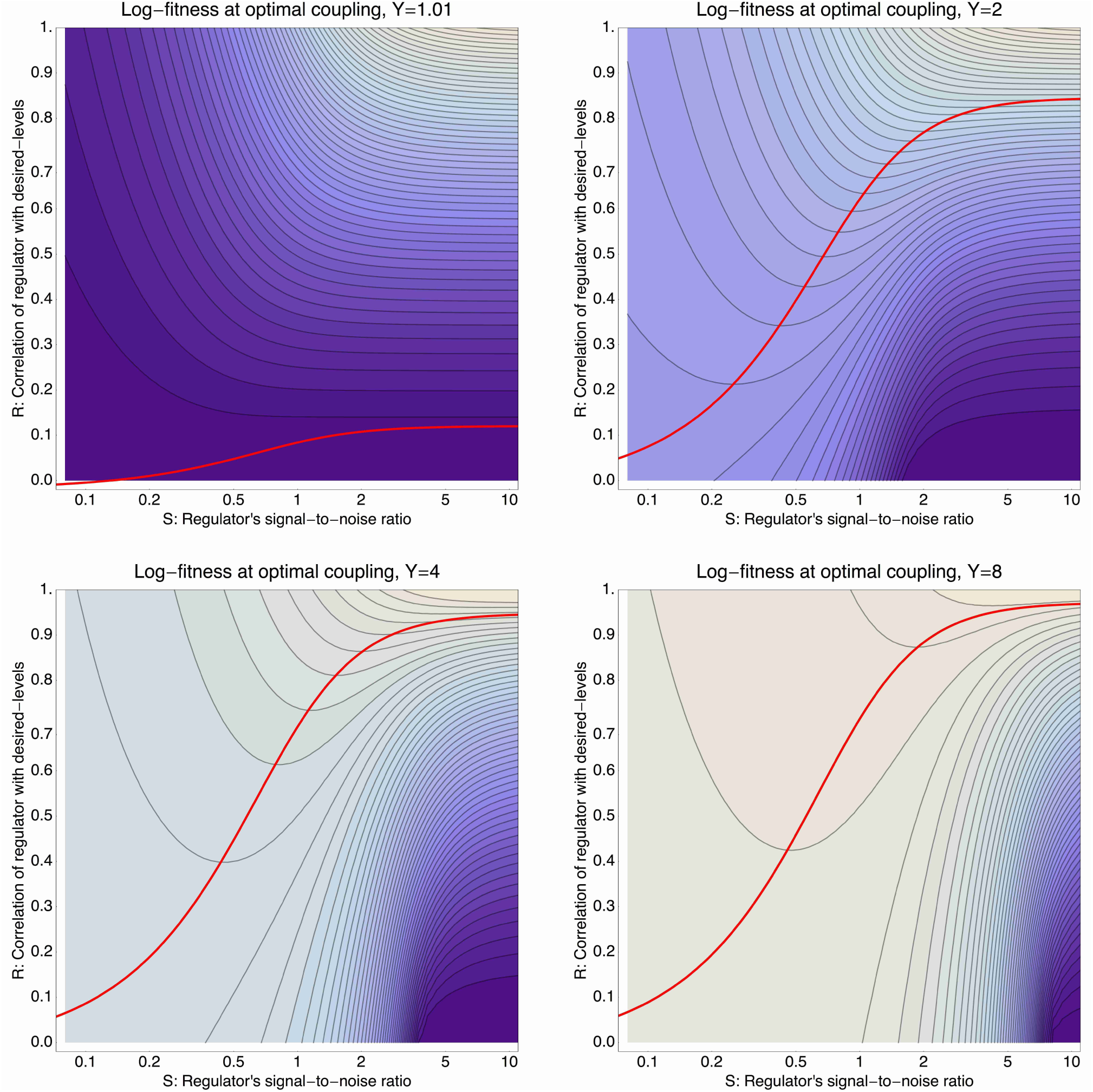
Log-fitness as a function of the signal-to-noise ratio *S* (horizontal axis) and correlation *R* of the regulator (vertical axis), for a promoter that is optimally coupled (*X* = *X*_∗_) to the regulator. The different panels correspond to log-fitnesses that are obtained for different values of the expression mismatch *Y* (indicated in the title of each panel). The contours run from −0.04 to −0.5 in the top-left panel, from −0.3 to −1.9 in the top-right panel, from −1 to −12 in the bottom-left panel, and from −2 to −30 in the bottom-right panel. The red curves show optimal signal-to-noise *S* as a function of the correlation *R*.

When the expression mismatch is small, i.e. *Y* = 1 corresponding to a variance in *µ_e_* that matches *σ*^2^ + *τ*^2^, then fitness generally increases with increasing *R* and *S*. However, the absolute value of the fitness increase is small. That is, for small *Y* promoters already have reasonably high fitness without regulation. As the value of *Y* increases, the log-fitness values start varying more dramatically as a function of *R* and *S*. Independent of the value of *Y*, the optimal fitness is always obtained for very high *R* and high *S*. However, when *Y* is large, an almost equally high fitness can be obtained by coupling to a ‘noisy’ regulator with low *S* and *R* = 0. In particular, when *Y* is large only regulators with very high correlation *R* and large signal-to-noise *S* can outcompete coupling to an entirely noisy regulator with *R* = 0 and small *S*. That is, in any situation where the desired expression levels vary significantly more than the width *σ* of the expression distribution, i.e. when *Y >* 1, promoters can substantially increase their fitness by coupling to a regulator that only acts to increase their noise. Moreover, to improve on this coupling to a ‘random’ regulator, a regulator has to be available with very high correlation *R* and large signal-to-noise. In other words, unless a regulator is available that very precisely regulates the promoter to attain its desired expression levels, best fitness can often be obtained by increasing the noise in the promoter's expression.

Note also that, whenever *Y* is larger than 1 and the correlation *R* is not very high, fitness generally decreases rapidly with the signal-to-noise of the regulator. That is, when a regulator has only moderate correlation with the desired expression levels of its target, low signal-to-noise is preferred. This suggests that regulators that are regulating targets whose desired expression levels correlate only moderately with the regulator's activity may be under selection for *lowering* their signal to noise ratios.

These considerations suggest different possible scenarios for the joint evolution of promoters and their regulators. On the one hand, when a regulator is coupled to a single promoter, or a set of promoters whose desired expression levels are perfectly correlated across environments, then a regulator can increase overall fitness by increasing the correlation *R* between the regulator's activities and the desired expression levels of its targets. In this way, regulation may evolve to become more precise over time. On the other hand, promoters often have an incentive to couple to regulators who only moderately correlate with their desired levels. Once a regulator is coupled to multiple promoters that have *different* desired expression levels, there is no way that the regulator can adapt its activities to correlate highly with the desired levels of all its targets, and such regulators will experience selection to become more noisy.

### Final noise levels under optimal coupling and signal-to-noise

We next consider what final noise levels 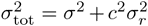 result when a promoter, with a certain expression mismatch *Y*, couples optimally to a regulator which has a certain correlation *R*, and whose signal-to-noise level has been optimized as well.

To this end we need to determine the jointly optimal coupling *X_∗_* and signal-to-noise *S_∗_* given a certain expression mismatch *Y* and correlation *R*. From equation (91) it is easy to see that fitness is maximized with respect to the signal-to-noise level *S* when

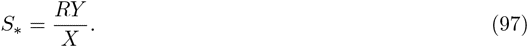

If we substitute this value back into the equation (91), we find that the optimal coupling *X_∗_* is now given by

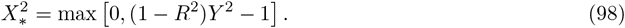

Note that, *Y*^2^ is the variation in desired expression levels, and *R*^2^*Y*^2^ is the amount of this variation that the promoter manages to ‘track’ when it is regulated by the regulator. Thus, (1 − *R*^2^)*Y*^2^ is precisely the remaining variance in desired expression levels that the promoter is unable to track. To bring this out more clearly, we substitute back our original parameters. We then find that when (1 *− R*^2^)*Y*^2^ *<* 1:

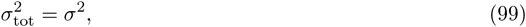

and when (1 − R2)Y 2 > 1

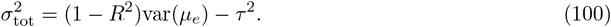

This brings out most clearly that, when the regulation is imprecise ((1 − *R*^2^)*Y*^2^ > 1), the final noise level that is evolved matches the fraction of the variance in desired expression levels var(*µ_e_*) that is not tracked by the regulation. In other words, the evolved transcriptional noise level of a promoter precisely reflects to what extent the promoter's regulation is not able to track the expression levels desired by the environment.

**Suppl. Fig. S8** shows the total noise level *σ*_tot_ as a function of *Y* and *R* when the promoter is optimally coupled to a regulator with optimal signal-to-noise. The figure illustrates that there are two regimes of solutions (‘phases’) separated by a phase boundary (thick black curve) that occurs at (1 − *R*^2^)*Y*^2^ = 1. On one side of this boundary, in the top-left of the figure, the final noise level *σ*_tot_ is essentially not different from the original noise level *σ*. This occurs either when *Y* < 1, i.e. when no regulation is necessary, or when very accurate regulation is available. Note that, similarly to what we saw in the last section, very high correlations *R* are necessary to realize this regime at larger values of *Y*. We call this the ‘basal noise regime’.

The largest part of the parameter space occurs on the other side of the phase boundary, which we call the ‘environment-driven noise regime’. Here the final noise level *σ*_tot_ becomes independent of the original noise *σ*, but is instead determined by the fraction of variation in desired expression levels var(*µ_e_*) that is not tracked by the regulation, i.e. by (1 − *R*^2^)var(*µ_e_*). The figure also indicates the optimal values of *S_∗_* as a function of *Y* and *R*. The optimal signal-to-noise *S_∗_* diverges at the phase boundary. That is, in the ‘basal noise regime’ regulators are preferred with signal-to-noise that is as high as possible. In contrast, for the majority of the parameter space in the environment driven noise regime signal-to-noise levels of 1 or less are preferred. That is, unless regulation is very precise, noisy regulators are typically preferred over precise regulators.

The figure also demonstrates that, unless *R* is close to one, the final noise increases with the variance in desired expression levels var(*µ_e_*) . Thus, unless there is a systematic correlation between the expression mismatch *Y* of a promoter, and the correlation of the regulator with highest available correlation, noise levels are expected to increase with the ‘plasticity’ var(*µ_e_*) that the environment requires of the promoters.

Similarly, the larger *Y*, the larger the remaining variance *Y′* = (1 *R*^2^)*Y* ^2^ tends to be after coupling to the regulator with the highest available correlation *R*. Whenever this remaining expression mismatch is *Y′* large, the promoter will have an incentive to couple to further regulators. That is, the theory also generally predicts that promoters with high var(*µ_e_*) tend to couple to more regulators.

Finally, we note that this theoretical model can easily be extended to the case of multiple promoters and regulators. In particular, because in our model the promoter's expression is a linear function of the regulatory inputs, the theory extends easily to promoters coupling to multiple regulators with different coupling constants. However, the regulatory network structure that will evolve in this general case will depend crucially on the correlation structure of the desired expression levels across all the promoters. Moreover, there might be many environmental changes that affect the optimal expression levels, but that cannot be sensed by any of the regulators, and this will constrain the extent to which regulators can optimize their activities to match the desired levels of their targets.

